# Deep sequencing artificially inflates estimates of microbial diversity

**DOI:** 10.64898/2026.09.01.748665

**Authors:** Lucas P. Henry, Eric Laderman, Joy Bergelson

**Affiliations:** Center for Genomics and Systems Biology, Dept. of Biology, New York University; Life Sciences, Simons Foundation

**Author notes:** **Correspondence:** (LPH), (JB).

## Abstract

Sequencing artifacts challenge accuracy and reproducibility when quantifying microbial diversity. To track error propagation in microbiome analyses, we analyze no-diversity amplicons, which are amplified from host genes with limited genetic diversity or from synthetic spike-ins. We find that sequencing at greater than 10^4^ reads exponentially increased no-diversity amplicon sequence variant (ASV) richness, with hundreds of ASVs observed per sample. This striking pattern was shared with microbial amplicons (16S rRNA, ITS, gyrB, rpoB), which revealed inflated Shannon diversity with an increase in read counts for both the community and within taxa. Comparing sequencing error profiles between no-diversity and microbial amplicons showed that truncating reads to shorter lengths and use of the AVITI Element platform can mitigate, but not abolish, the impacts of artificial inflation; we recommend caution when read depths vary orders of magnitude between samples. Overall, utilizing no-diversity amplicons can help optimize parameters to improve estimates of microbial diversity.

## INTRODUCTION

With the revolution of high-throughput sequencing, we can now use amplicon profiling to illuminate the previously unseen microbial world with remarkable ease. This has allowed us to discover that the microbiome contributes to many aspects of host health and potentially offers novel solutions to problems in medicine, agriculture, and conservation^1–3^. However, the biological complexity of the microbiome continues to pose significant challenges due to the vast number and diversity of microbial cells within a typical sample^4^. Moreover, many technical factors are known to bias estimates of microbial diversity, which reduces accuracy and reproducibility in microbiome studies^5–9^. If we aim to use the microbiome to improve host health, then we need to improve methods that accurately and reproducibly measure microbial diversity.

The common recommendation for accurately assessing the enormous complexity of the microbiome is to sequence more deeply. Indeed, a strong predictor of microbial diversity is sampling effort^10–17;^ the more reads sequenced, the more taxa discovered. This effect is believed to be a consequence of the power gained in detecting rare taxa. However, the reality is more complex. Instead, biases in microbial diversity can emerge because samples were sequenced at different depths^14,16,17^. This problem is exacerbated by the ease of generating hundreds of millions of reads, given decreasing sequencing costs and the widespread adoption of amplicon sequence variants (ASVs)^18^. For example, high-quality sequencing reads with a PHRED score of 30 represents an error rate of 0.1% per base, which could propagate error substantially at the rate of 25 million errors for 250 bp reads in 100 million reads. For over a decade, the potential consequences of read depth has been a contentious area of vigorous debate in microbial ecology^13–17,19^. Yet, it is common for read depth to vary orders of magnitude across samples within a study, and this rarely results from true biological variation. As such, there is great need to develop approaches that detect errors and mitigate the consequences on microbial diversity estimates.

Here, we propose using “no-diversity” amplicons to evaluate errors in microbial diversity estimates. No-diversity amplicons are amplified from synthetic spike-ins or host housekeeping genes that are normally used to estimate microbial load^20,21^. Because spike-ins and housekeeping genes have extremely limited genetic diversity and are co-amplified during library preparation, no-diversity amplicons can serve as a powerful indicator of the conversion of raw data into microbial diversity estimates. Using this approach, we discovered a surprising inflation in diversity at depths exceeding 10^4^ reads/sample in both no-diversity and microbial amplicons. We then explore how tuning parameters during ASV calling can mitigate this inflation and show that the AVITI Element removes erroneous ASVs more efficiently than Illumina Novaseq. Together, our analyses build a robust framework to optimize parameters to more accurately estimate microbial diversity.

## RESULTS

### No-diversity amplicons exhibit high and variable diversity

We first focus on the *GIGANTEA* (GI) amplicon, which is a single-copy host gene that was previously developed to estimate microbial load in *Arabidopsis thaliana*^20,22^. There is genetic diversity in GI between natural accessions^23^, but because *A. thaliana* is diploid, there can only be a maximum of two alleles per individual. Thus, we expect a maximum of two ASVs, but no ASV diversity per individual. The data were generated from wild *A. thaliana* collected in Michigan^24^ and in New York City, NY. Many of the libraries were sequenced twice across different Illumina Novaseq runs, and GI ASVs were called from all runs using the default DADA2 parameters implemented in QIIME2^25^. Reads were truncated to 250 bp for ASV calling, as sequence quality dropped in the last 50 bp (Supp. Fig. 1).

GI richness varied across hundreds of samples and seven sequencing runs, with a mean 83 GI ASVs per sample and as high as 609 ASVs observed in one sample (Fig. 1A). The mean and variance in GI ASV richness significantly differed across sequencing runs (mean: Kruskall Wallis X^2^ = 1359.7, df = 6, p < 2.2e-16; variance: Levene’s F_6,2205_ = 437.83, p < 2.2e-16). GI Shannon diversity was also significantly different for mean and variance across runs (Fig. 1B; mean: Kruskall Wallis X^2^ = 1374.0, df = 6, p < 2.2e-16; variance: Levene’s F_6,2205_ = 316.24, p < 2.2e-16). We also included a synthetic 16S rRNA spike-in in Run07 as another no-diversity amplicon, which exhibited significantly higher inflation than the GI ASV (Supp. Fig. 2).

**Fig. 1:**
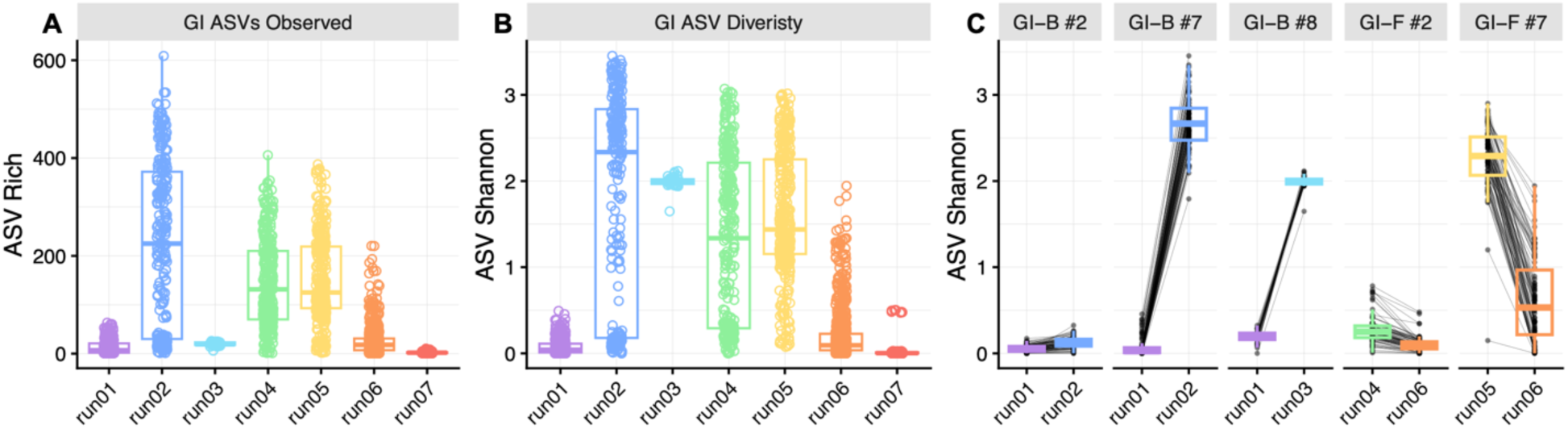
GI amplicons show variation in ASV calling across sequencing runs. Points represent individual *A. thaliana* and colors show sequencing runs. A) As GI is a no-diversity amplicon, the number of ASVs observed should be low, but we observed as high as 609 GI ASVs in a single individual. B) Shannon diversity for GI should be ∼0, but we observed a range in diversity as well. C) Representative plots of GI ASV diversity for samples that were sequenced twice. Black lines connect GI Shannon diversity from individuals across the sequencing runs. Facet labels “GI-B” reflect when co-amplified with bacterial 16S rRNA and ”GI-F” reflect when co-amplified with fungal ITS1-2. The change in GI Shannon diversity differs within pools and between sequencing runs.

This inconsistency in GI diversity did not result from library preparation, as variation was observed from the same samples between runs (Fig. 1C); the dominant ASV remained consistent (Supp. Fig. 3). Rather, inconsistencies were driven by variation in read depth across sequencing runs, with higher depths per sequencing pool generating higher richness (Supp. Fig. 4). However, GI ASV richness varied from tens to hundreds even at 1000 reads/sample (Supp. Fig. 5). Read depth alone is not sufficient to explain these patterns and highlights the value of no-diversity amplicons to expose technical artifacts.

### Case study: No diversity-amplicon helps fix an egregious artificial inflation of diversity

We first present an important example of using no-diversity amplicons to improve estimates of microbial diversity. In Run03, we observed a marked increase in GI ASV Shannon diversity (Fig. 1C). Further inspection revealed 11 GI ASVs above 1% relative abundance, while only a single GI ASV was above 1% relative abundance for the same samples in Run01 (Fig. 2A). This diversity in Run03 occurred because SNPs accumulated (Fig. 2B). No sequence variation was observed within the first 210 bp, while 13 SNPs occur in the last 40 bp (Supp. Fig. 6). Strikingly, the top 5 most abundant bacterial taxa also show an accumulation of SNPs in a similar location (Fig. 2B); informative sequence variation beyond family or genus level assignments is not expected for 16S rRNA amplicons^26,27^. While it is possible that these SNPs reflect biologically meaningful variation, the shared regions in microbial data that match the problematic GI variation suggest these are the result of technical artifacts.

**Fig. 2:**
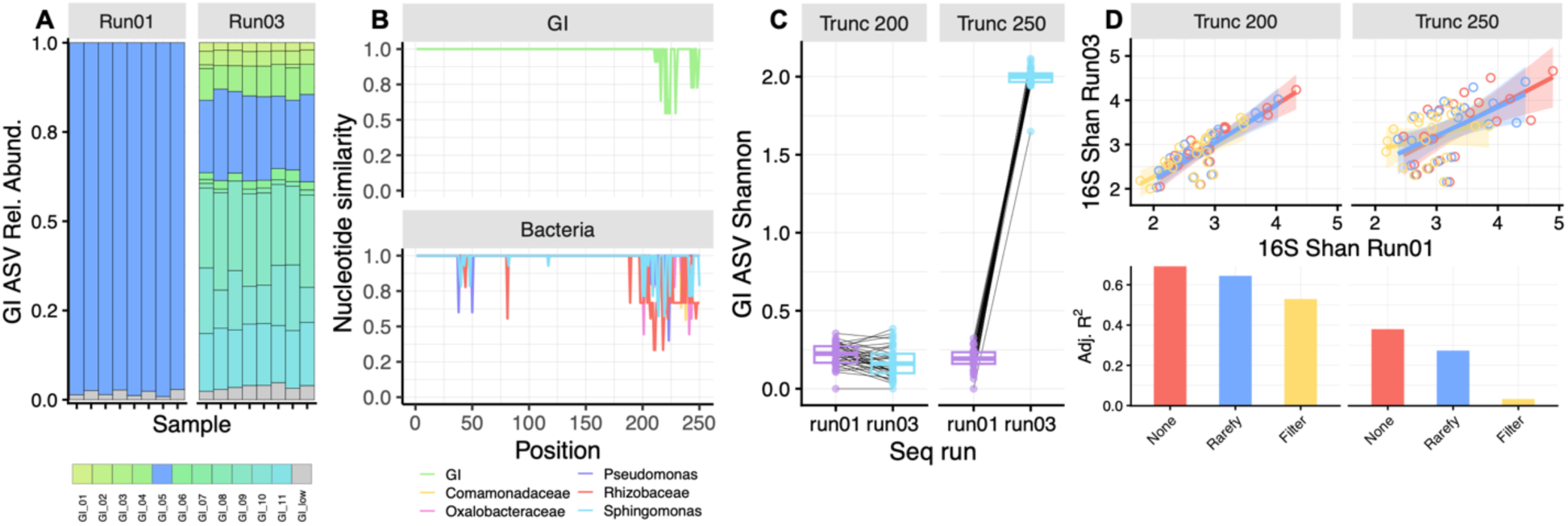
A case study for using no-diversity amplicons to correct an artificial inflation of diversity. A) Representative stacked barplots showing an increased diversity of GI ASVs from the same eight samples in Run01 and Run03. Colors represent different ASVs, and grey represents all <1% relative abundance ASVs. B) Comparing nucleotide similarity (1.0 = identical nucleotides across ASVs) at each position along the GI ASVs (green line) and top 5 bacterial taxa ASVs (colored lines) from Run03. C) The difference in GI ASV diversity between sequencing runs is reduced when ASVs are truncated to 200 bp compared to 250 bp. Lines connect the sample between sequencing runs. D) Correlation in Shannon diversity between the two sequencing runs for the two truncation lengths. Each point is a sample, colored by pre-processing as either none (red), rarefied (blue), or filtered for ASVs <= 0.5% (yellow). Bottom shows adjusted R^2^ values for the correlation between sequencing runs.

We reasoned that truncating to 200 bp would remove this problematic region of the ASVs and increase similarity between the two runs. Indeed, the difference in mean GI Shannon diversity at 200 bp was reduced compared to 250 bp (Fig. 2C, Supp. Table 1). The 200 bp truncation for 16S rRNA amplicons also showed a stronger correlation of microbial Shannon diversity between the two runs (Fig. 2D, Supp. Table 2). The adjusted R^2^ was 0.69 for 200 bp ASVs, but only 0.39 for 250 bp ASVs. One reason for this difference could be uneven read depth among samples between the two truncation lengths^16^. To test this, the data was filtered (removing <0.5% relative abundance ASVs) or rarefied (depth = 500 reads/sample, subsampled 100 times). The 200 bp ASVs still outperformed the 250 bp ASVs with the highest adjusted R^2^, and neither rarefaction nor filtering improved the correlation compared to no pre-processing (Fig. 2D). Together, these results demonstrate a clear benefit of using no-diversity amplicons to improve estimates of microbial diversity.

### Read depth and truncation length expose idiosyncrasies in ASV calling

We next expanded our analysis to other common amplicons that characterize microbial communities. This includes fungi (ITS1-2) and two other bacterial genes, gyrB and rpoB, which evolve faster and provide finer resolution than the 16S rRNA amplicon^28,29^; only ASVs assigned to phylum-level classification were included in analyses. We then leveraged the insights gained from the Run03 analyses to identify two generalizable factors that impact estimates of ASV diversity: read depth and truncation length.

First, ASV richness tended to increase exponentially with read depth for all amplicons (Fig. 3A). The magnitude of inflation varied across amplicons, with ITS reaching a maximum 241 ASVs in an individual sample, while 16S rRNA reached its maximum at 1606 ASVs, gyrB at over 7000 ASVs, and rpoB at 2404 ASVs. While it is common to see the positive correlation between depth and ASV richness^10,11,13,14,16^, it is surprising to see its nearly exponential increases.

**Fig. 3:**
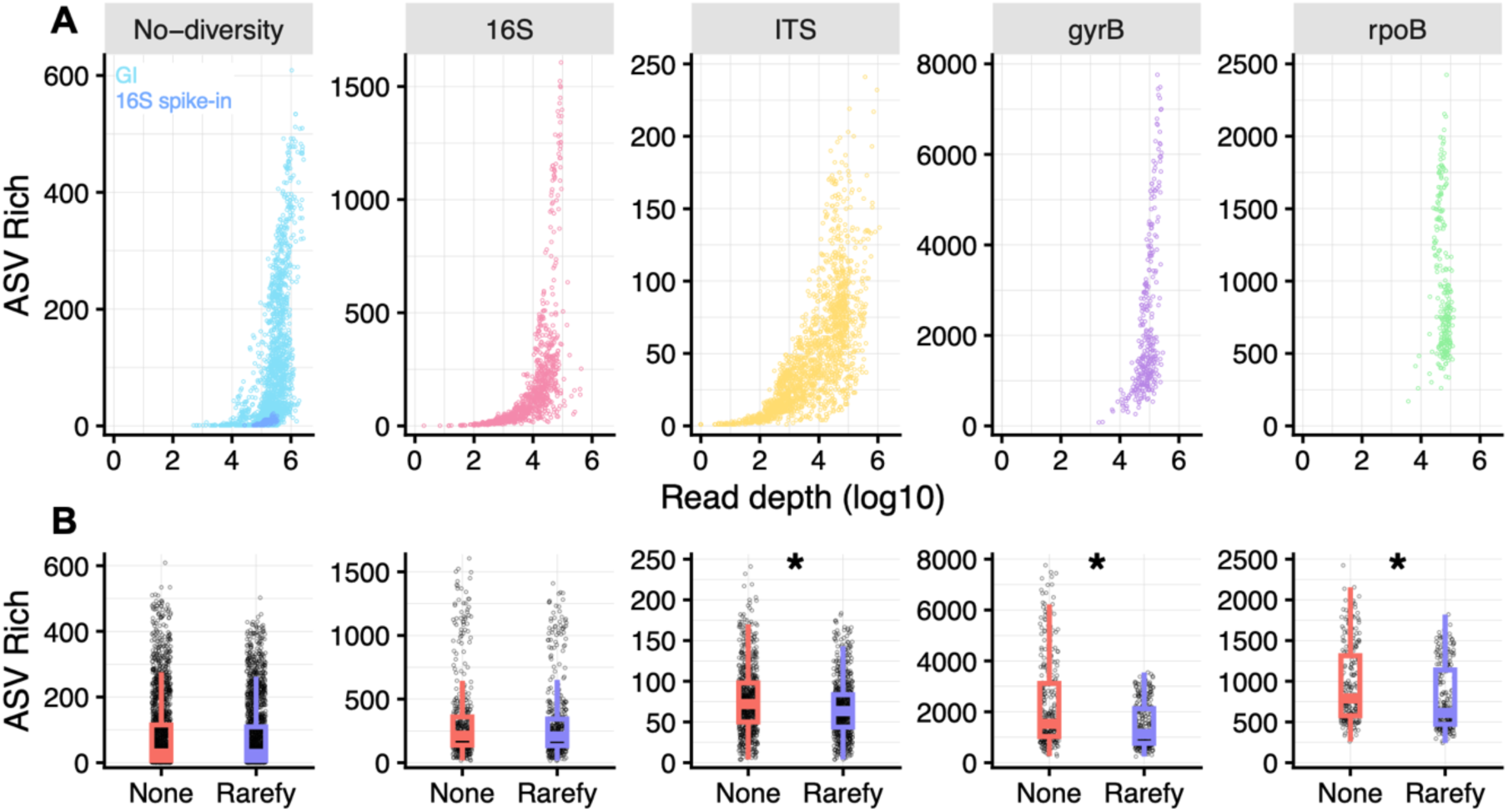
ASV richness increases exponentially above 10^4^ reads across all amplicons surveyed. A) Each point represents the ASV richness per individual sample at 250 bp truncation length. No-diversity includes both GI (light blue) and the 16S spike-in (dark blue). B) Rarefying at 10,000 reads/sample reduces ASV richness, but only significantly for ITS, gyrB, and rpoB amplicons (asterisk represents p < 0.001).

This exponential increase likely emerges during ASV calling in the DADA2 algorithm^30^. DADA2 first builds an error model based on quality scores and then sorts sequences into tentative ASVs based on abundance. If an ASV is more abundant than expected from error rates alone, then it is a true ASV. Noisy ASVs are those that are rare and differ in a few base pairs relative to a more abundant ASV. For example, singleton ASVs are deemed noise because the error-learning model cannot differentiate these from errors; singletons are removed in default settings. Increasing sequencing depth, however, distorts the balance between noise and abundance because errors will be numerically more abundant, resulting in artificially inflated diversity. To some extent, rarefaction can reduce ASV richness (Fig. 3B, Supp. Table 3), but neither rarefaction nor 0.5% relative abundance filtering mitigated the exponential inflation above 10^4^ reads in no-diversity and microbial amplicons (Supp. Fig. 7). This suggests consistent technical artifacts that cannot be resolved by filtering or rarefying alone.

Second, we tested the impact of the interaction between truncation length and read depth on ASV diversity. We first pooled together reads from all samples per sequencing run to focus on factors that underlie likely technical artifacts for each amplicon. Then, we subsampled to different read depths (1x10^3^, 1x10^4^, 5x10^4^, 1x10^5^, and 5x10^5^ reads; 10 replicates per depth) to mimic a common range of read depths^16^ at six different truncation lengths (150, 175, 200, 225, 250, 275 bp).

For all amplicons, higher read depths are associated with higher diversity, especially above 10^4^ reads (Fig. 4A; Supp. Table 4); similar patterns were observed for richness (Supp. Fig. 8, Supp. Table 5). However, the impact of truncation length and read depth on Shannon diversity differed by amplicon (interaction Wald X^2^ = 531.08, df=25, p < 0.0001, Supp. Table 4). For example, the 16S rRNA and the co-amplified GI both showed an increase in Shannon diversity for read lengths of 200 bp or longer, but ITS and co-amplified GI idiosyncratically varied. The high resolution amplicons, gyrB and rpoB, displayed consistent Shannon diversity values across truncation lengths, with the exception of gyrB at 275 bp, where the majority of reads were filtered out in the DADA2 algorithm. Across all amplicons, the 275 bp truncation length retained the fewest reads (Supp. Fig. 9).

**Fig. 4:**
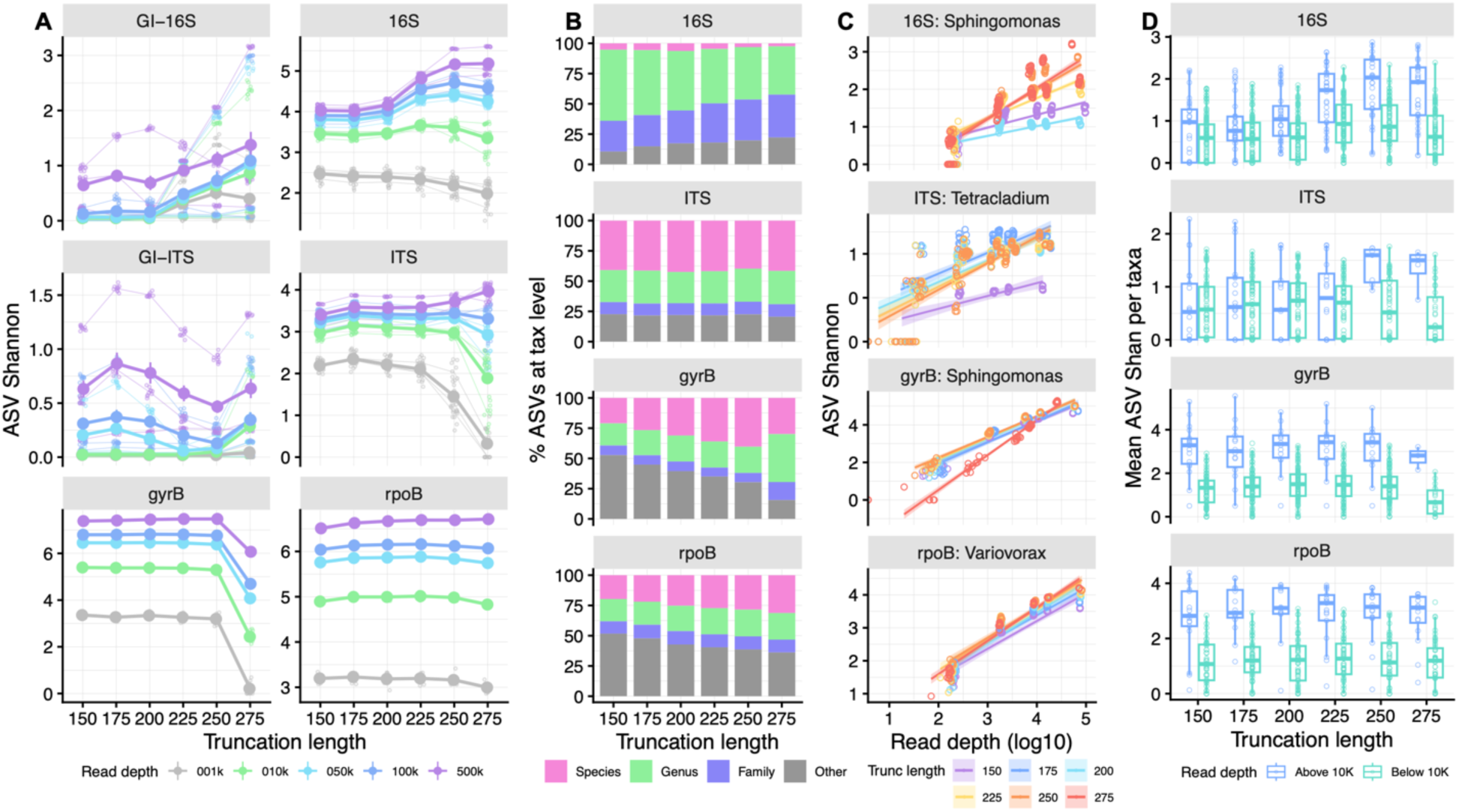
Truncation length and read depth impact ASV Shannon diversity in different ways across amplicons. Thick lines and points represent mean ± standard error across all sequencing runs per sequencing depth; note gyrB and rpoB were only on one run. Thin lines show the different runs, and small points represent the subsampled pools (10 replicates/read depth). A) Relationship between truncation length and read depth varies across amplicons. B) Percentage ASVs at different taxonomic levels for each truncation length, faceted by microbial amplicons. C) Shannon diversity within the top taxa is correlated with total read depth for a given sample, but the slope depends on truncation length. D) Read depth systematically impacts Shannon diversity within taxa both above and below 10^4^ reads. Each point represents the mean Shannon diversity for each taxon at the different truncation lengths.

We next focus on truncation length alone. An important truncation tradeoff exists: too short may remove informative sequences for taxonomic resolution, while too long can result in both increased noise from sequencing and/or increased loss of reads during quality filtering. We observed two dynamics between the standard and high-resolution amplicons (Fig. 4B). For 16S, most ASVs were assigned to genus, but longer amplicons decreased resolution in taxonomic assignments. ITS ASVs were instead robust to amplicon length. For gyrB and rpoB, longer amplicons increased the percentage of ASVs assigned at the species level, with the exception of 275 bp for gyrB where substantial read loss was observed (Supp. Fig. 9). Thus, the appropriate truncation length will depend on amplicon and desired resolution of taxonomic assignment.

These technical factors have important ramifications. First, read depth impacts not just the discovery of new ASVs, but also genera and species (Supp. Fig. 10, Supp. Table 6). Given the exponential increase of ASV richness at greater than 10^4^ read depths, we reasoned that this is also likely to impact diversity estimates within dominant taxa. Indeed, for the most abundant taxa assigned at the genus level, ASV Shannon diversity within each taxon was positively correlated with read depth (Fig. 4C, Supp. Table 7); a similar relationship was observed across the top five most abundant taxa per amplicon (Supp. Fig. 11). Second, the slope of the read depth-diversity relationship depended on the truncation length of the amplicon (interaction Wald X^2^ = 11.81, df = 3, p = 0.008), and this interaction impacted 16S and ITS more than gyrB and rpoB (Fig. 4C).

Because read depth increases ASV diversity within taxa, total community diversity may be shaped by individual taxa with greater than 10^4^ reads/taxa within the microbiome. To test this, we focused on the subsampling at 10^5^ reads per replicate. We classified taxa present in at least 50% of replicates as abundant (≧10^4^ reads) or rare (<10^4^, but ≧10^3^ reads); all taxa with less than 10^3^ reads were excluded. Then, ASV richness and Shannon diversity were calculated per taxa for each replicate. We used genus-level assignments for 16S and ITS, but species-level assignments for gyrB and rpoB, given the increased resolution for these amplicons^28,29^. Note that both abundant and rare groups contained a mix of taxonomic levels (Supp. Fig. 12).

Indeed, taxa at greater than 10^4^ reads had higher ASV richness (Supp. Fig. 13, Supp. Table 8), with mean ASV richness approximately twice as high for 16S and ITS and 7-10 times higher for gyrB and rpoB. This further impacted Shannon diversity, with systematically higher diversity in abundant amplicons (Fig. 4D, Supp. Table 9). Truncation length further exacerbated the differences between amplicons for the abundant taxa at longer truncation lengths (interaction Wald X^2^ = 19.70, df = 1, p < 0.0001). This analysis suggests that there is systematic increase in ASV richness and diversity for taxa above 10^4^ reads, which in turn can disproportionately inflate diversity estimates.

One possible fix is to increase the sensitivity to maximum expected errors based on quality scores (maxEE). maxEE models the error distribution of PHRED scores in Illumina sequencing and varies with each nucleotide within a read, as opposed to a more simplistic mean filtering of quality scores. We compared the effect of default maxEE = 2 in QIIME2^25^ and stricter cutoffs at 1 and 0.1 on ASV calling. Only the strictest cutoff of maxEE = 0.1 substantially reduced ASV richness, but this was achieved through reducing read depth by at least 50% (Supp. Fig. 14).

This mitigated the 10^4^ inflation of richness and diversity (Supp. Fig. 15), but the severe reduction in read depth resulted in increasingly noisy estimates between replicates (Supp. Fig. 16, Supp. Table 10). Strict maxEE filters can minimize ASV inflation, but is an unrealistic solution given the extreme read loss and increase in noise across samples.

### AVITI sequencing mitigates technical artifacts better than Illumina sequencing

The above analyses explored the effects of technical artifacts on ASV diversity estimates, guided by our expectations for no-diversity amplicons. To explore other solutions, we next compared the impact of technical artifacts with the relatively new AVITI Element sequencing platform.

The AVITI platform uses avidity chemistry that employs rolling circle amplification and high affinity, multivalent bonding with nucleotide fluorophores to improve quality during base calling^31^. Indeed, the average PHRED scores on AVITI were higher than on Illumina (Supp. Fig. 17), but dips in PHRED scores were more randomly dispersed. PHRED scores are also fully continuous in AVITI data, but binned into four groups from the Illumina Novaseq, which alters the variance in quality scores along the read (Supp. Fig. 17). These differences may impact DADA2 performance, which was developed for Illumina data^30^. To compare platforms, identical samples were sequenced and analyzed as above. These analyses focus on wild *A. thaliana* in New York City, NY, and includes soil microbiomes, which are more diverse than plant microbiomes^32^, to investigate the impact of these technical artifacts on high diversity microbiomes.

Using the subsampled depths at six truncation lengths, AVITI showed improved performance over Illumina in two key ways. First, AVITI data were more robust to technical artifacts for no-diversity amplicons, with lower ASV richness in deep sequencing and no difference between 200 and 250 bp truncation lengths (Fig. 5A, Supp. Table 11); Novaseq inflation was 2.5-5X higher than AVITI inflation at 5x10^5^ reads. Second, across all amplicons, AVITI data purged erroneous ASVs at approximately twice the rate of Illumina during maxEE filtering (Fig. 5B, Supp. Table 12; see Supp. Fig. 18 for all data). Even with these improvements, we observed diversity inflation above 10^4^ reads (Supp. Fig. 19). Beyond the 200 and 250 bp comparison in Figs. 5A-B, truncation length also impacted diversity, and the magnitude and direction of the difference between AVITI and Novaseq depended on the amplicon, with larger impacts on 16S and ITS amplicons (Supp. Fig. 20, Supp. Table 13).

**Fig. 5:**
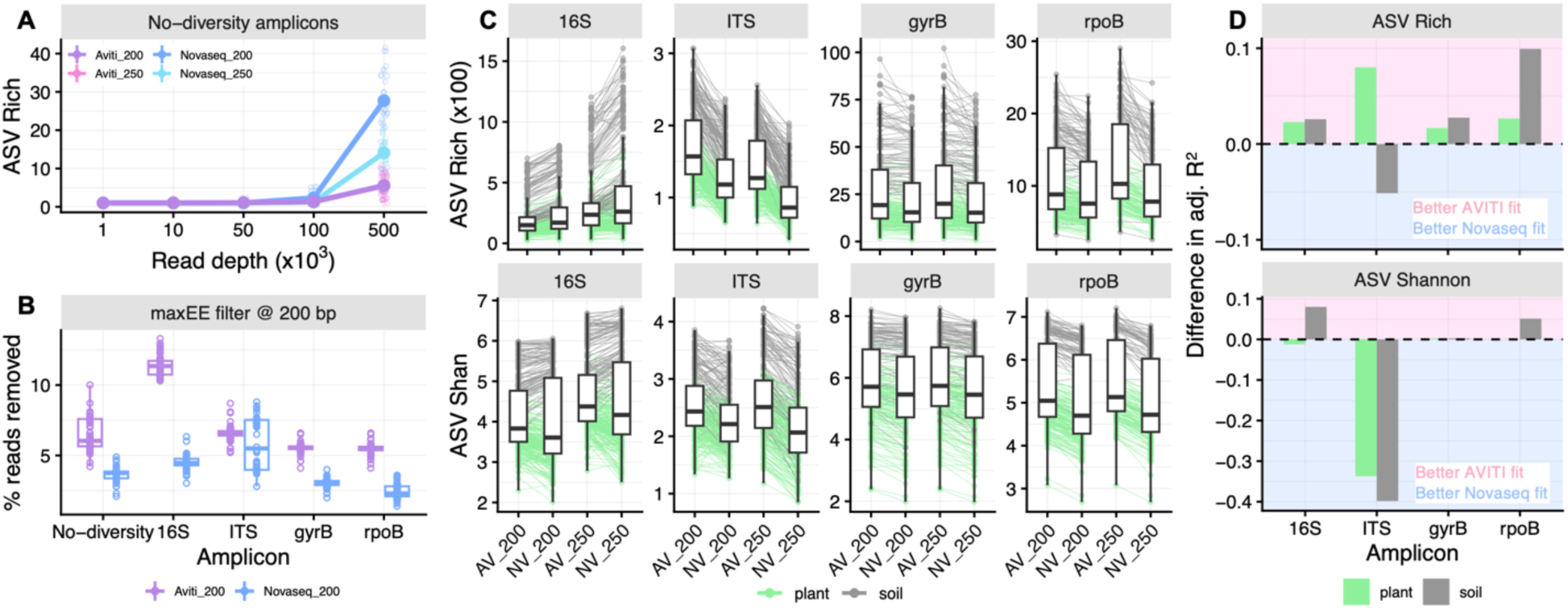
AVITI sequencing improves estimates of microbial diversity. A) No-diversity ASV richness is inflated at 5x10^5^ read depth when using the Novaseq platform. B) For all amplicons, more reads are removed during the maxEE filtering step from data generated on the AVITI platform. C) Alpha-diversity differences at 200 or 250 bp for both platforms (AV = AVITI, NV = Novaseq). Richness (scaled by 100X) and Shannon diversity are per sample, and lines connect the samples across both truncation lengths, colored by whether the sample was plant or soil. D) Difference in the adjusted R^2^ for the correlation between 200bp and 250bp truncation length within each platform as a measure of robust microbial diversity estimates. Bars are colored by plant or soil, while plot area is shaded by whether the adjusted R^2^ was higher in the AVITI (pink) or Novaseq (blue) data.

Finally, we compared microbial diversity estimates from true biological samples. For this analysis, we matched the read depth per sample and focused on 200 and 250 bp truncation lengths. Diversity estimates across all amplicons were sensitive to interactions between sequencing platform and truncation length (Fig. 5C, Supp. Table 14). ASV richness for 16S amplicons were inflated in Novaseq data, particularly for the higher-diversity, soil microbiomes at 250 bp. In the other amplicons, richness and diversity was generally higher on the AVITI. To test if AVITI data is more robust to truncation length as a proxy for error accumulation, we examined the fit for the correlation between truncation lengths with each platform. The fit was high for nearly all comparisons, with adjusted R^2^ > 0.9 (Supp. Table 15). In particular, AVITI data displayed better fits for high diversity soil microbiomes than Novaseq (Fig. 5D). The one exception with a better fit in Novaseq data was ITS, but inconsistencies in taxonomic assignment between truncation lengths and correlations were poor for both platforms (Supp. Fig. 21). Taken together, these results suggest that the AVITI platform improves estimates of microbiome diversity.

## DISCUSSION

Our analysis across amplicons yields important generalities. First, deep sequencing (>10^4^ reads) artificially inflates diversity, as observed in no-diversity and microbial ASVs. Second, shorter ASVs minimize the artificial inflation, but optimizing truncation length will depend on the amplicon. Third, these technical artifacts impact diversity estimates, particularly as more abundant taxa display higher rates of ASV inflation than less abundant taxa. These issues will be especially exacerbated in datasets where samples range in read depth by orders of magnitude, as these effects emerge at 10^4^ reads. Taken together, we offer three general recommendations to improve estimates of microbial diversity: 1) avoid richness as a diversity metric, 2) truncate to shorter amplicons to minimize sequencing errors, and 3) the AVITI Element platform reduces technical artifacts compared to the Illumina Novaseq.

Richness should be avoided as a diversity metric because it was substantially impacted by all technical factors we examined. Shannon diversity was more robust, with smaller changes from technical artifacts. The most troubling phenomenon we observed was the 2-10X increased richness for abundant taxa above 10^4^ reads (Supp. Fig. 13) and corresponding increase in Shannon diversity (Fig. 4D). We have not seen this issue documented in the literature, nor can we invoke any plausible biological mechanism. The concerning consequence of this phenomenon would be if a manipulation enriches a dominant taxa (e.g., microbial blooms^33^) compared to a control group, which could skew effects of the manipulation based simply on read abundances. For 16S and ITS, shorter amplicons reduced the within-taxa inflation, but otherwise it is a difficult problem to rectify if read depths vary substantially between samples.

Identifying the correct truncation length has been a perennial problem for microbiome analysis. The general guidance is to qualitatively assess PHRED profiles to remove enough data to capture noise, but not too much. Instead, no-diversity amplicons enable a precise, quantitative assessment of truncation length, as it is crystal clear where erroneous nucleotides appear (Fig. 2B). Furthermore, perhaps counterintuitively, shorter amplicons had better taxonomic resolution for 16S rRNA (Fig. 4B). There is a growing movement to use full-length 16S rRNA to characterize the microbiome at higher resolution^34–36^, but our results suggest caution. As the full-length 16S approach uses a range of sequencing platforms, it will be necessary to evaluate error profiles through no-diversity amplicons to control for artificial inflation of sequencing errors in each platform, as we did (Figure 5).

In conclusion, no-diversity amplicons serve as a powerful diagnostic approach to detect and mitigate technical artifacts in estimating microbial diversity. Diversity of the microbiome is susceptible to many technical factors that may distort inferences^6^, but carefully designed analyses can minimize these effects. Our approach adds another tool with which to do so and control the consequences of read depth variation^13–17,19^. Ultimately, ecological diversity is an important measure and the starting point in investigating host-microbiome interactions^37^–our contribution here is to help reproducibly measure microbial diversity to leverage the potential benefits of the microbiome for improving host health.

## ACKNOWLEDGEMENTS

We thank the Bergelson lab for helpful feedback. We acknowledge the Zegar Family Foundation for their generous support of the NYU Genomics Core. LPH was supported by the Charles H. Revson Foundation. This work was funded by grant SFI-PD-Grant-01308072 of the Simons Foundation to JB.

## DATA AND CODE AVAILABILITY

Sequencing data for the Michigan dataset is available uploaded to NCBI SRA, as Bioprojects PRJNA1279102 for 16S rRNA and PRJNA1282642 for ITS1-2; both Bioprojects include the GI amplicons that were co-amplified with the 16S and ITS amplicons. NYC data will be made available upon publication. Code will be made available upon publication.

## ONLINE METHODS

The data analyzed here comes from two surveys of microbiome variation in wild *Arabidopsis thaliana* populations. The first dataset is a survey of *A. thaliana* populations across different land-use types in Michigan^24^. The second dataset surveyed a single *A. thaliana* population in New York City, NY. The amplicon libraries were prepared differently between the two datasets, but the majority of our comparisons focus on within dataset comparisons. Libraries were generated per sample once but then were generally sequenced twice per amplicon across different sequencing runs (Supp. Table 16). This approach enables comparisons that identify robust parameters to estimate microbial diversity, while controlling for variation that results from library preparation.

## AMPLICON LIBRARY PREPARATION

There were two differences in the sample processing between the two datasets, the DNA extraction method and PCR cycle number during library preparation. Our inferences are robust to the differences in sample processing, as we performed our analyses within each dataset and uncovered similar patterns between the datasets.

In the DNA extraction, the Michigan samples were initially ground with ∼0.5 ml of garnet rocks or a mix of silica beads (bead size ranged from 0.5 to 3.0 mm) under liquid nitrogen. DNA was then extracted using a modified home-brew method^24^. DNA was normalized to ∼10 ng/µl prior to library prep; if the DNA concentration was below 10 ng/µl, it was not diluted. For the NYC samples, samples were ground with a mix of 1mm silica beads and 4mm glass beads under liquid nitrogen. DNA was extracted using the Zymo Quick DNA/RNA MagBead kit following manufacturer’s directions. DNA concentrations ranged from 1-15 ng/µl; no normalization was performed prior to library preparation.

Libraries were prepared using a two-step, dual indexed PCR procedure that amplifies the genes of interest in the first PCR and adds Illumina adapters in the second PCR. For the Michigan dataset, 10 cycles were used in the first PCR and 30 cycles in the second PCR. Samples were treated with ExoSAP to remove unannealed primers between first and second PCR. For the NYC dataset, 15 cycles were used in the first PCR and 25 cycles in the second PCR; no ExoSAP treatment was performed. Libraries were prepared using OneTaq HotStart polymerase (NEB) for the Michigan dataset and Q5 HotStart polymerase (NEB) for the NYC dataset.

We amplified a total of five amplicons across the datasets analyzed (Supp. Table 17): GI, 16S V5V6V7, ITS1-2, gyrB, and rpoB. gyrB and rpoB amplicons were produced only for the NYC dataset. We adjusted annealing temperatures accordingly for each amplicon (Supp. Table 17).

The GI and 16S spike-in served as the no-diversity amplicons. All GI amplicons were co-amplified with either 16S or ITS following the host-associated microbe PCR protocol^20^. The 16S spike-in sequence was used only in the NYC dataset and designed with the same 16S V5V6V7 primer sites, but with random nucleotides in between the primer sites. The synthetic spike-in was the same length as the 16S amplicon. The spike-in was synthesized by Twist Biosciences, cloned into pUC18 plasmid, and introduced into E. coli DH5a cells. Plasmids were then isolated using the Zyppy Plasmid Miniprep kit (Zymo Research Corporation). Plasmids were diluted to 1x10^-4^ ng/ul for the first PCR and only added to soil samples during library preparation.

Libraries were pooled per PCR plate (∼90 samples/plate) using equal volumes and cleaned using SPRI beads. After SPRI bead cleaning, libraries were further concentrated and extraneous bands were removed using the Zymoclean Gel DNA Recovery kit. Quality of pools was assessed using TapeStation D1000 reagents, and quantity was assessed using Qubit reagents. PCR plate pools were normalized to the same molarity prior to sequencing.

All sequencing was configured as dual-indexed, single-end 300 bp reads. Michigan and NYC samples were sequenced on the Illumina Novaseq 6000 platform, performed by the NYU Genomics Core. For Novaseq data, the reads were basecalled using Picard IlluminaBasecallsToFastq version 2.23.8^38^, with APPLY_EAMSS_FILTER set to false. Reads were demultiplexed using barcode_splitter^39^. Only NYC samples were sequenced on the AVITI Element, performed by AUGenomics (San Diego, CA). For AVITI, the Cloudbreak Freestyle High kit was used to circularize libraries for avidity chemistry, and the reads were basecalled and demultiplexed using Bases2Fastq.

## BIOINFORMATICS

After demultiplexing, Illumina adapters and polyG sequences were removed using fastp^40^; reads with fewer than 250bp were removed. Samples were uniquely labeled with i5/i7 indices, but not each amplicon. Each amplicon was then split using the following tools. GI, 16S, and ITS reads were split using the primer sequence as input into BBtools BBDuk^41^. The 16S spike-in was then filtered from the 16S dataset through an additional round of BBDuk using the first 50bp of the synthetic sequence. Because gyrB and rpoB primers contain many degenerate sites, seqfu^42^ was used to filter these amplicons.

Because our initial analyses showed the importance of read depth in shaping ASV diversity, we subsampled to five different read depths that encompass a common range in data sets: 1x10^3^, 1x10^4^, 5x10^4^, 1x10^5^, and 5x10^5^ reads, with 10 replicates per read depth. Data was subsampled per amplicon and sequencing run to control for run-to-run variation. To do this, we first generated a base pool of ∼5.5x10^6^ reads per run and amplicon to increase the chances that the 10 replicates of the 5x10^5^ subsampling were sampling distinct reads. To make this base pool, we sampled each individual sample at equal depths. For example, if a run had 500 samples, we randomly sampled 11,000 reads per sample and concatenated them together into the base pool. From this base pool, we subsampled at the different read depths to generate the 10 replicates. All subsampling was performed using BBtools reformat^41^.

We also subsampled using BBTools format for the comparisons of individual samples that were sequenced on both AVITI and Novaseq platforms. Read depth was first calculated for each individual sample, and then we subsampled down to the lower of the two to match read depth.

All ASVs were called using QIIME2 v2024.10. First, primer sequences were removed using Cutadapt^43^. Then, DADA2^30^ was used to call ASVs. ASVs were called per PCR plate pool, which enabled us to examine how any previous processing (as these were batched for DNA extraction and library preparation) potentially impacted ASV calling. To examine the effects of different truncation lengths, the “trunc-len” command was set to the six different lengths (150, 175, 200, 225, 250, 275 bp); no left-trim was performed. Most analyses used the default maxEE = 2 as the quality filtering parameter. To increase the stringency of this quality filter, for the Novaseq subsampled data, we ran subsequent analyses at maxEE = 1 and 0.1 to compare performance. After ASV calling, taxonomy was assigned using the classify-sklearn Bayes Classifier^26^, except for ITS. ITS ASVs were classified using BLAST^44^ and ClassifyITS^45^.

Appropriate reference taxonomic databases were used for each amplicon. The GI reference database included the *GIGANTEA* sequence from Arabidopsis as well as *GIGANTEA* from *Capsicum annuum* and *E. coli* 16S sequence to insure specificity. The Greengenes database^26,46^ was used for 16S sequences, trimmed to the V5V6V7 region. ITS sequences were classified using the Unite database^47^. We used a custom gyrB database to classify sequences^48^, which was built using ∼55,000 gyrB sequences compiled from publicly available data from JGI and NCBI. The rpoB ASVs were classified using the FROGS rpoB database^29^. For all analyses, only ASVs assigned to at least phylum were used; any ASV with only a kingdom assignment was removed from analyses. We did not perform any other filtering, unless explicitly described. Data then was imported into phyloseq^49^ to calculate ASV richness (i.e., the number of unique ASVs per sample) and Shannon diversity for the various analyses.

To compare the effects of filtering and rarefaction, we removed ASVs with less than 0.5% relative abundance per sample. To rarefy, ASV tables were subsampled without replacement 20 times to 10^4^ reads/sample, and then the mean ASV richness and Shannon diversity was calculated from those 20 rounds of subsampling.

To understand the effects of the 10^4^ reads threshold for diversity inflation, we used our subsampled data for each amplicon at the read depth of 1x10^5^ reads. This read depth was used so that several taxa could be detected at greater than 10^4^ reads. We first used the “tax_glom” in phyloseq to sum all ASVs at either the genus level for 16S and ITS or the species level for gyrB and rpoB. We note that this included ASVs that could not be resolved at this taxon level (e.g., Comamonadaceae family at the Genus level in the 16S data). A taxon was considered abundant if total reads were greater than 10^4^. We only included taxa that were present in at least 5/10 of the replicates, and we excluded any taxa with less than 10^3^ reads (i.e. <1% relative abundance). Mean ASV richness and Shannon diversity was then calculated for each of these taxa individually in the ten replicates and compared across the six truncation lengths.

## STATISTICAL ANALYSES

Our general approach was to first use non-parametric tests to assess general factors that impacted estimates of microbial diversity for the data shown in Figures 1-3. Then, to focus on specific parameters, we used linear mixed models in data shown in Figures 4-5 and the associated supplemental figures. We used random effects to control for general factors like sequencing run–that is, to ask whether these correlations are robust when accounting for the random effect. All analyses were performed in R. Linear mixed models were implemented using lmer^50^. Normality of residuals was checked using DHARMa package^51^. Significance of interactions was tested with the appropriate sum of squares (e.g., Type II or Type III) using car^52^. We next walk through each set of analyses in detail.

We initially tested for the differences in ASV richness and diversity for GI amplicons (data shown in Fig. 1) using non-parametric Kruskal-Wallis tests to provide an overview of differences between the eight different sequencing runs. We also tested for unequal variances using Levene’s test.

In the case study (data shown in Fig. 2), our approach to find parameters that increased reproducibility was to compare diversity between Run-01 and Run-03. We performed a linear regression that asked if the diversity estimates (ASV richness or Shannon diversity) from Run01 predicted the diversity estimates in Run03. We examined this correlation for the 200 and 250 bp truncation lengths for each of the pre-processing approaches: no filtering, rarefaction to 10^4^ reads, or filtering out ASVs <= 0.5% relative abundance.

To examine the impact of rarefaction on ASV richness (data shown in Fig. 3B), we performed non-parametric Kruskal-Wallis tests separately for each amplicon.

To examine the interactions between truncation length and read depth on ASV Shannon diversity, we fitted the following mode [1], with the alpha-diversity measure as either Shannon diversity or log-10 transferred ASV richness. We used the subsampled data at the five different sequencing depths and different truncation lengths. The approach here allows for us to ask whether amplicons differ in the interaction between read depth and truncation length. The significance of the interactions was evaluated using Type III Wald X^2^ tests.

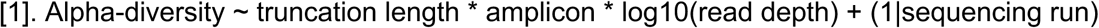

To test if the discovery of taxa at different levels (e.g., species-level assignment or ASV) was shaped by truncation length and read depth, we fitted the following model [2]. We used the subsampled data at the five different sequencing depths and different truncation lengths. We used amplicon as a random effect here because we wanted to ask if this relationship was generally predicted from interactions between read depth, truncation length, and taxa-level. The significance of the interactions was evaluated using Type III Wald X^2^ tests.

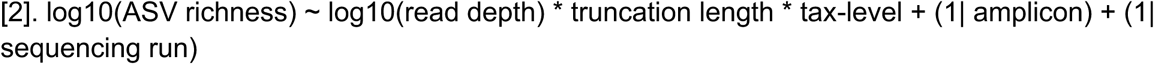

To test if the ASV Shannon diversity of the most abundant taxa was impacted by read depth and truncation length, we fitted the model below [3]. We used the subsampled data at the five different sequencing depths and different truncation lengths. The three-way interaction allows us to test if the amplicon (e.g., 16S, ITS, etc) differs in this relationship. Shannon diversity was square-root transformed to ensure normality of residuals. The significance of the interactions was evaluated using Type III Wald X^2^ tests.

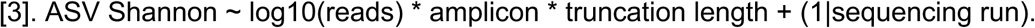

To test if taxa that were abundant above 10^4^ reads also were more diverse, we fitted the model below [4]. The abundance group is either greater than 10^4^ reads or <10^4^ but greater than 10^3^ reads from the subsampled data to 10^5^ reads/replicate. We fitted two different models using log-10 transformed mean ASV richness per taxa or square root-transformed mean Shannon diversity per taxa. We used amplicon as a random effect here to identify if the effects of truncation length and abundance group were robust across amplicons. The significance of interactions was evaluated using Type II Wald X^2^ tests.

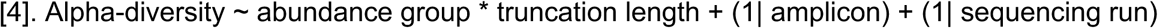

To test how varying maxEE impacted the noise in Shannon diversity estimates, our response variable was standard deviation. We fitted the model below [5]. We used the subsampled data at the five different sequencing depths and different truncation lengths. The standard deviation of Shannon diversity was square root-transformed to ensure normality of residuals. Amplicon was used as a random effect here to identify if the interactions were robust across amplicons. The significance of interactions was assessed using Type III Wald X^2^ tests.

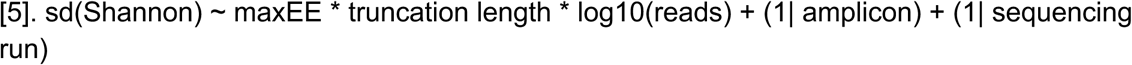

To test for the effects of the sequencing platform (Novaseq versus AVITI) on alpha-diversity estimates, we fitted the model below [6]. We used the subsampled data at the five different sequencing depths and different truncation lengths. We fitted two different models using log-10 transformed mean ASV richness per taxa or square root-transformed mean Shannon diversity. We used a four-way interaction and assessed significance using Type III Wald X^2^ tests. Our random effect here was sample type (plant versus soil), which allowed us to test if these effects were robust across plant and higher-diversity soil samples. We confirmed that the model was not overfit by comparing AIC and BIC values with a less complex model that did not include the four-way interaction.

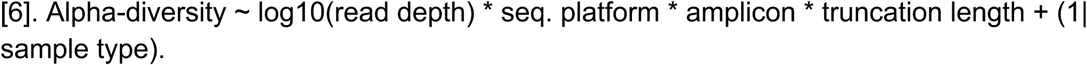

To test for the effects of the sequencing platform on read filtering during ASV calling, we fitted the model below [7]. We used the subsampled data at the five different sequencing depths and different truncation lengths. Our approach was to focus on how the sequencing platform impacted read filtering depending on the amplicon. Here, we used truncation length and sample type as random effects to determine if this relationship was robust across these parameters. Significance was assessed using Type II Wald X^2^ tests.

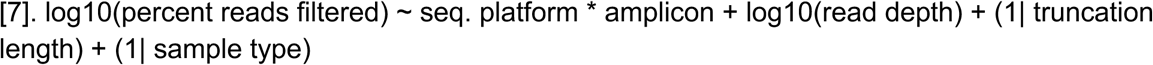

To test if the sequencing platform impacted the percentage of the microbiome resolved to meaningful taxonomic identity (e.g., beyond phylum or class level) and if this differed by amplicon, we fitted the model below [8]. We used the subsampled data at the five different sequencing depths and different truncation lengths. Our random effect approach here was to ask if this was robust across read depth, sample type (plant versus soil) and truncation length. Significance was assessed using Type II Wald X^2^ tests.

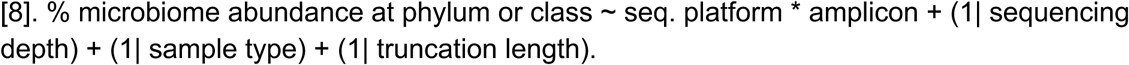

To test the impact of the sequencing platform on diversity estimates for real samples, we used individual samples that were subsampled to the same read depth and fitted the model below [9]. We then compared the effects of two different truncation lengths (200, 250 bp) based on our insights from the case study. Here, we included sample type as a main effect because we were interested in understanding if these technical artifacts differentially impacted plant and high-diversity soil samples. We first compared the correlation in ASV richness and Shannon diversity for samples between the truncation lengths within sequencing platform by each sample type. To do so, we filtered each dataset separately and performed linear regression as [9]:

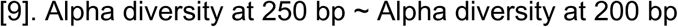

This enabled us to use the adjusted R^2^ for each fit to understand how error propagation (i.e., longer reads are more likely to have more errors) differed between low (plant) and high (soil) diversity microbiomes for each sequencing platform.

Next, we fitted two different models using log-10 transformed mean ASV richness per taxa or square root-transformed mean Shannon diversity. We initially fitted a four-way interaction between sequencing platform, sample type, amplicon, and truncation length, but this four-way interaction was not statistically significant. So, we refitted the model with the two significant three-way interactions as shown below [10]. We assessed significance using Type III Wald X^2^ tests. We confirmed that the model was not overfit by comparing AIC and BIC values with a less complex model that did not include the two three-way interactions. The random effect used here was each plate pool, which reflects the ∼90 samples that were processed together from DNA extraction through library prep and initially ASV calling.

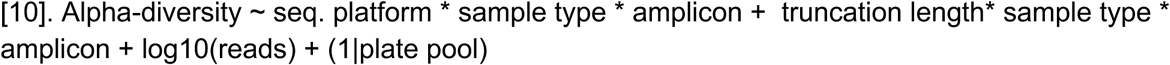

## Notes

### Competing Interest Statement

The authors have declared no competing interest.

