## Supplemental Figures and Tables for "Deep sequencing artificially inflates estimates of microbial diversity"

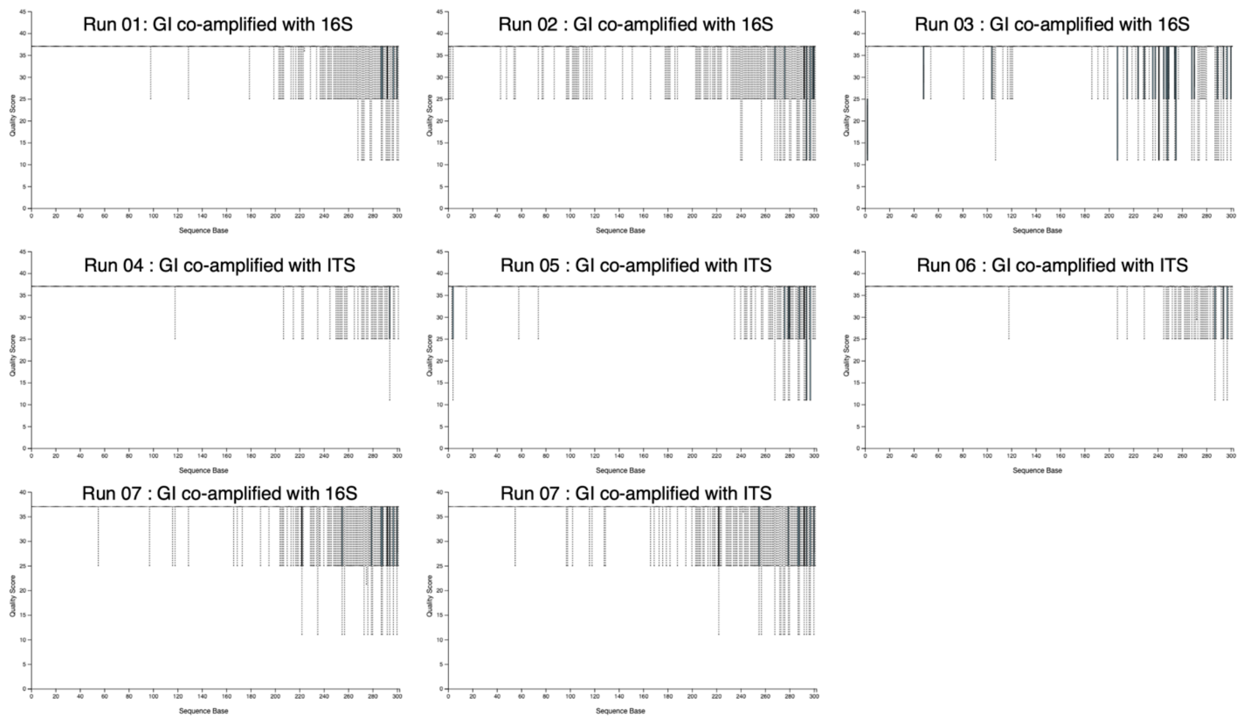


**Supp Fig 1:** Example traces for PHRED quality scores for GI sequences across different sequencing runs. Plots were generated using QIIIME2 demux visualization and are generated from the same library pools that were sequenced across the different platforms. Note that Run 1 and Run 3 plots are associated with the dataset that is the focus for the section “Case study: No diversity-amplicon helps fix an egregious artificial inflation of diversity”.

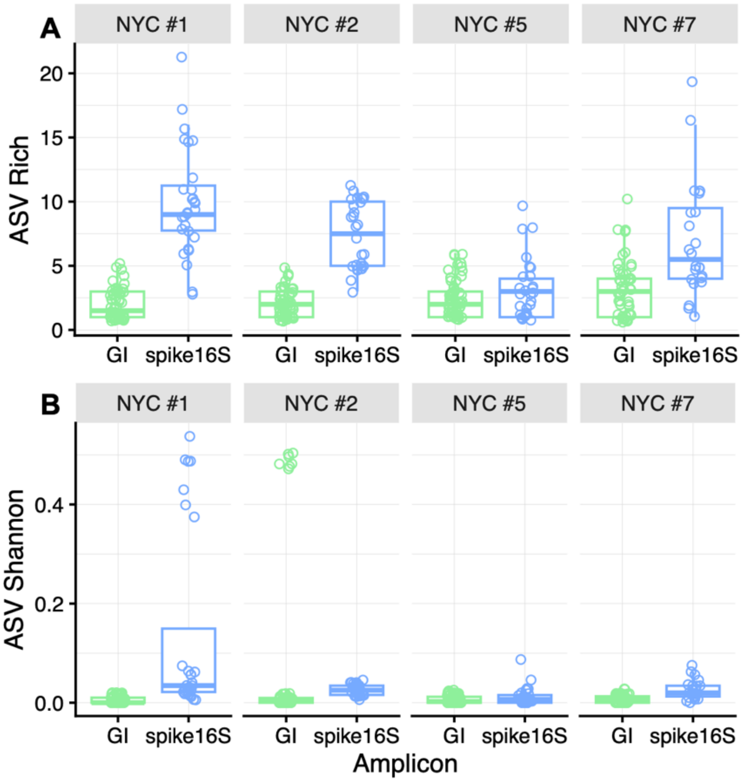


**Supp Fig 2:** 16S spike-ins also show inflations in ASV richness and Shannon, though are more exaggerated compared to the GI ASV. Each point represents an individual sample. For 16S these are soil samples, and GI are *A. thaliana* plants. A) ASV richness is above 0 for both 16S spike and GI, but the 16S spike is significantly higher than GI (Kruskal-Wallis $X^2$ = 105.1, df = 1, p < 2.2e-16). B) ASV Shannon diversity is also significantly higher in the 16S spike than GI (Kruskal-Wallis $X^2$ = 78.40, df = 1, p < 2.2e-16).

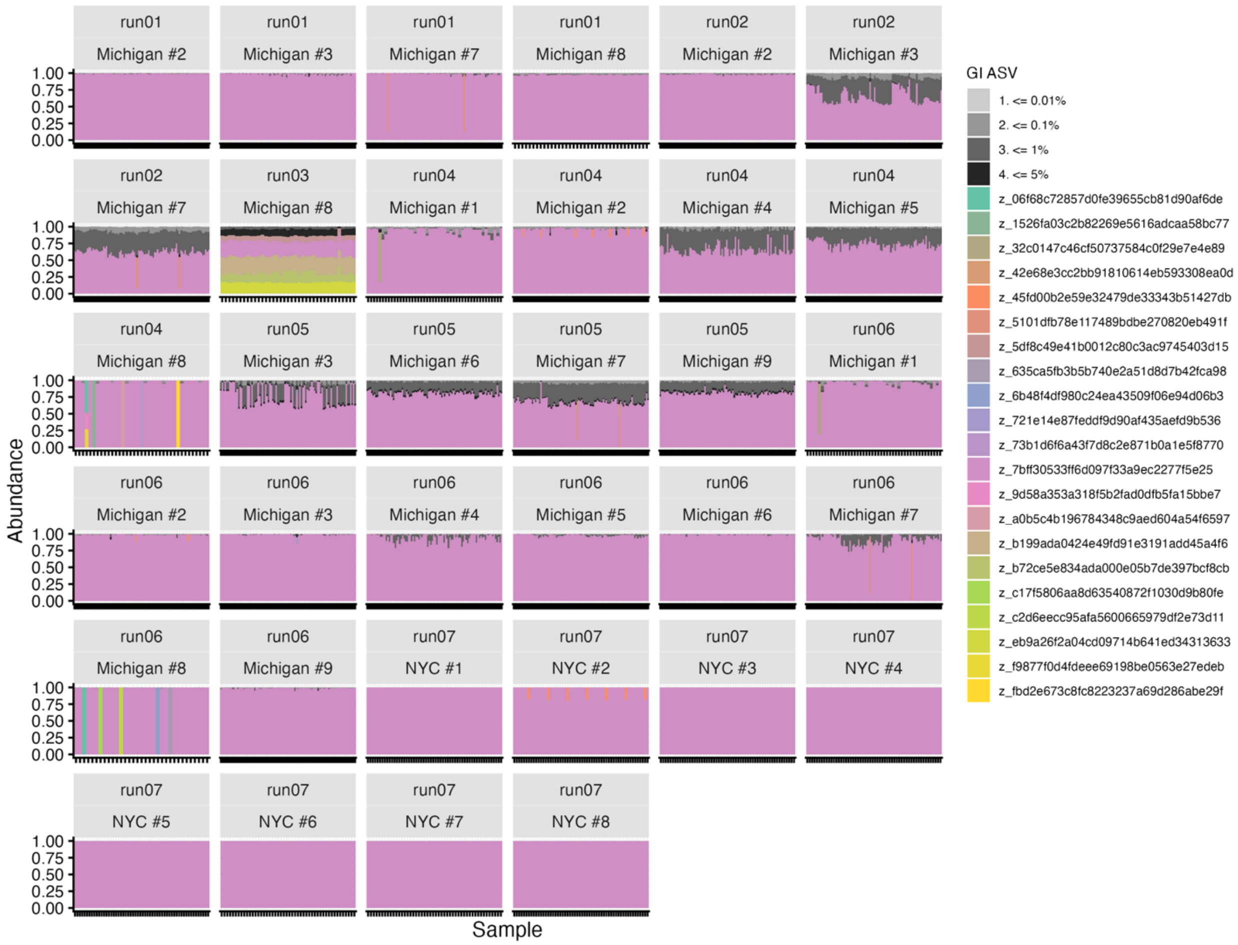


**Supp Fig 3:** GI ASV relative abundance distributions across the seven runs, faceted by library prep plate and sequencing run. Sample names are presented to ease visualization. Ticks on the x-axis denote individual samples. Note that many ASVs are grouped together within each level at <=5% relative abundance.

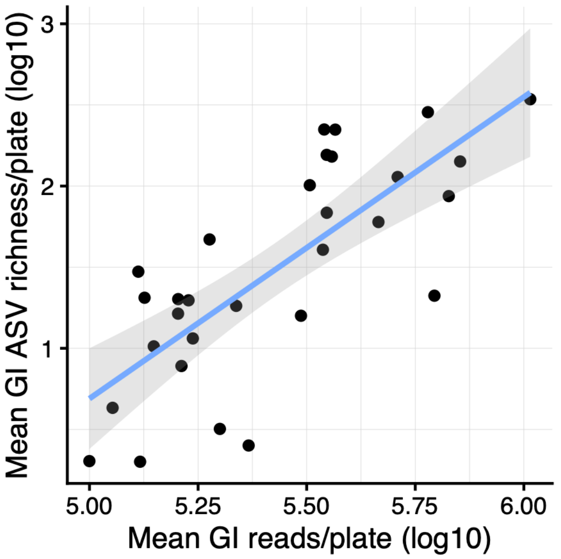


**Supp Fig 4:** Mean GI ASV richness is positively correlated with read depth (b = 1.35 +/- 0.35 standard error, df = 24.82, t = 3.84, p = 0.0008, marginal $R^2$ = 0.36). Each point represents a library pool, which was ~90 individuals processed together as a plate through DNA extraction, library prep, and data analyses. The correlation was evaluated using a mixed linear effects model, with DNA plate and sequencing run as random effects.

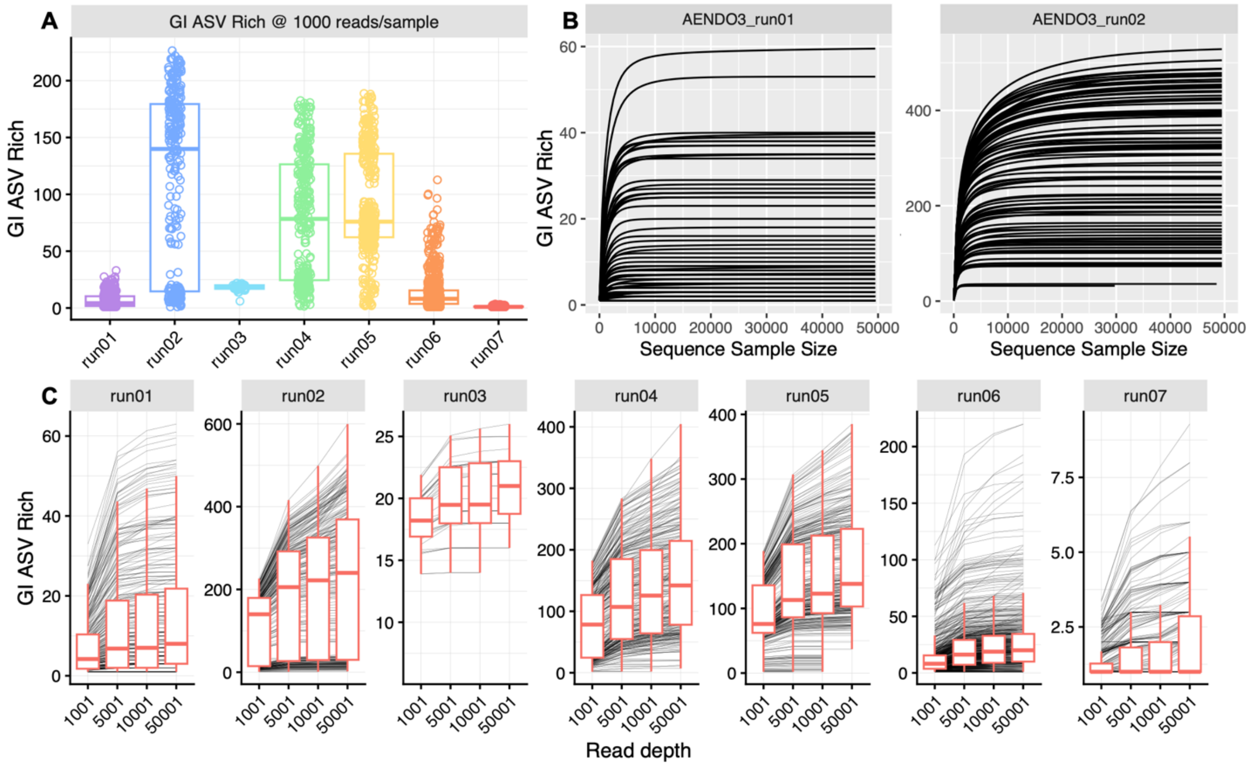


**Supp Fig 5:** Variation in read depth alone does not explain the artificial inflation of GI ASV richness. A) GI ASV richness at 1000 reads per sample, which demonstrated that read depth alone does explain the range of ASV richness observed. B) Example rarefaction curves for one plate that was sequenced on Run01 and Run02. Note the differences in the y-axis for the range of ASV richness, highlighting that library prep alone is not responsible for these differences. X-axis was trimmed to 50000 reads for visualization purposes, as most samples had higher than 50000 reads. C) Differences in GI ASV richness observed at rarefied to at different read depths. Boxplots show each run, and thin individual lines show individuals across the different depths.

```
GI_05      AGAGTATGGAGCTGGGATTGACTCGGCAATTAGTCATACGCGCCGAATTTTGGCAATCCT 60
GI_07      AGAGTATGGAGCTGGGATTGACTCGGCAATTAGTCATACGCGCCGAATTTTGGCAATCCT 60
GI_09      AGAGTATGGAGCTGGGATTGACTCGGCAATTAGTCATACGCGCCGAATTTTGGCAATCCT 60
GI_06      AGAGTATGGAGCTGGGATTGACTCGGCAATTAGTCATACGCGCCGAATTTTGGCAATCCT 60
GI_11      AGAGTATGGAGCTGGGATTGACTCGGCAATTAGTCATACGCGCCGAATTTTGGCAATCCT 60
GI_02      AGAGTATGGAGCTGGGATTGACTCGGCAATTAGTCATACGCGCCGAATTTTGGCAATCCT 60
GI_08      AGAGTATGGAGCTGGGATTGACTCGGCAATTAGTCATACGCGCCGAATTTTGGCAATCCT 60
GI_03      AGAGTATGGAGCTGGGATTGACTCGGCAATTAGTCATACGCGCCGAATTTTGGCAATCCT 60
GI_01      AGAGTATGGAGCTGGGATTGACTCGGCAATTAGTCATACGCGCCGAATTTTGGCAATCCT 60
GI_04      AGAGTATGGAGCTGGGATTGACTCGGCAATTAGTCATACGCGCCGAATTTTGGCAATCCT 60
GI_10      AGAGTATGGAGCTGGGATTGACTCGGCAATTAGTCATACGCGCCGAATTTTGGCAATCCT 60
           ************************************************************

GI_05      AGAGGCACTCTTTTCATTAAAACCATCTTCTGTGGGGACTCCATGGAGTTACAGTTCTAG 120
GI_07      AGAGGCACTCTTTTCATTAAAACCATCTTCTGTGGGGACTCCATGGAGTTACAGTTCTAG 120
GI_09      AGAGGCACTCTTTTCATTAAAACCATCTTCTGTGGGGACTCCATGGAGTTACAGTTCTAG 120
GI_06      AGAGGCACTCTTTTCATTAAAACCATCTTCTGTGGGGACTCCATGGAGTTACAGTTCTAG 120
GI_11      AGAGGCACTCTTTTCATTAAAACCATCTTCTGTGGGGACTCCATGGAGTTACAGTTCTAG 120
GI_02      AGAGGCACTCTTTTCATTAAAACCATCTTCTGTGGGGACTCCATGGAGTTACAGTTCTAG 120
GI_08      AGAGGCACTCTTTTCATTAAAACCATCTTCTGTGGGGACTCCATGGAGTTACAGTTCTAG 120
GI_03      AGAGGCACTCTTTTCATTAAAACCATCTTCTGTGGGGACTCCATGGAGTTACAGTTCTAG 120
GI_01      AGAGGCACTCTTTTCATTAAAACCATCTTCTGTGGGGACTCCATGGAGTTACAGTTCTAG 120
GI_04      AGAGGCACTCTTTTCATTAAAACCATCTTCTGTGGGGACTCCATGGAGTTACAGTTCTAG 120
GI_10      AGAGGCACTCTTTTCATTAAAACCATCTTCTGTGGGGACTCCATGGAGTTACAGTTCTAG 120
           ************************************************************

GI_05      TGAGATAGTTGCTGCGGCCATGGTTGCAGCTCATATTTCCGAACTGTTCAGACGTTCAAA 180
GI_07      TGAGATAGTTGCTGCGGCCATGGTTGCAGCTCATATTTCCGAACTGTTCAGACGTTCAAA 180
GI_09      TGAGATAGTTGCTGCGGCCATGGTTGCAGCTCATATTTCCGAACTGTTCAGACGTTCAAA 180
GI_06      TGAGATAGTTGCTGCGGCCATGGTTGCAGCTCATATTTCCGAACTGTTCAGACGTTCAAA 180
GI_11      TGAGATAGTTGCTGCGGCCATGGTTGCAGCTCATATTTCCGAACTGTTCAGACGTTCAAA 180
GI_02      TGAGATAGTTGCTGCGGCCATGGTTGCAGCTCATATTTCCGAACTGTTCAGACGTTCAAA 180
GI_08      TGAGATAGTTGCTGCGGCCATGGTTGCAGCTCATATTTCCGAACTGTTCAGACGTTCAAA 180
GI_03      TGAGATAGTTGCTGCGGCCATGGTTGCAGCTCATATTTCCGAACTGTTCAGACGTTCAAA 180
GI_01      TGAGATAGTTGCTGCGGCCATGGTTGCAGCTCATATTTCCGAACTGTTCAGACGTTCAAA 180
GI_04      TGAGATAGTTGCTGCGGCCATGGTTGCAGCTCATATTTCCGAACTGTTCAGACGTTCAAA 180
GI_10      TGAGATAGTTGCTGCGGCCATGGTTGCAGCTCATATTTCCGAACTGTTCAGACGTTCAAA 180
           ************************************************************

GI_05      GGCCTTGACGCATGCATTGTCTGGGTTGATGAGATGTAAGTGGGATAAGGAAATTCATAAAAGAGCATCA 250
GI_07      GGCCTTGACGCATGCATTGTCTGGGTTGATGAGATGTAAGTGGGATAAGTAAATTCATAAAATATCAAAA 250
GI_09      GGCCTTGACGCATGCATTGTCTGGGTTGATGAGATGTAAGTGGGATAAGGAAATTCATAAAATATCATCA 250
GI_06      GGCCTTGACGCATGCATTGTCTGGGTTGATGAGATGTAAGTGGGATAAGTAAATTCATAAAATATCAACA 250
GI_11      GGCCTTGACGCATGCATTGTCTGGGTTGATGAGATGTAAGTGGGATAAGTAAATTCATAAAATATCATCA 250
GI_02      GGCCTTGACGCATGCATTGTCTGGGTTGATGAGATGTAATTTGTATAATTAAATTCATAAAAGAGCATCA 250
GI_08      GGCCTTGACGCATGCATTGTCTGGGTTGATGAGATGTAATTTTTATAATTAAATTCATAAAAGAGCATCA 250
GI_03      GGCCTTGACGCATGCATTGTCTGGGTTGATGAGATTTAATTTTTATAATTAAATTCATAAAATATCAAAA 250
GI_01      GGCCTTGACGCATGCATTGTCTGGGTTGATTATATTTAATTTTTATAATTAAATTCATAAAATATCATCA 250
GI_04      GGCCTTGACGCATGCATTGTCTGGGTTGATGAGATTTAATTTTTATAATTAAATTCATAAAATATCATCA 250
GI_10      GGCCTTGACGCATGCATTGTCTGGGTTGATGAGATTTAATTTGTATAATTAAATTCATAAAATATCATCA 250
           ****************************** * ** *** *   ****  ************ * **  *
```

**Supp Fig 6:** Clustal-W alignment of the 11 variable GI ASVs above 1% relative abundance detected in Run03. Sequences are identical until 210 bp, but then 13 SNPs accumulate in the last 40 bp of the 250 bp amplicon. Stars denote identical sequences, while gaps show the location of SNPs.

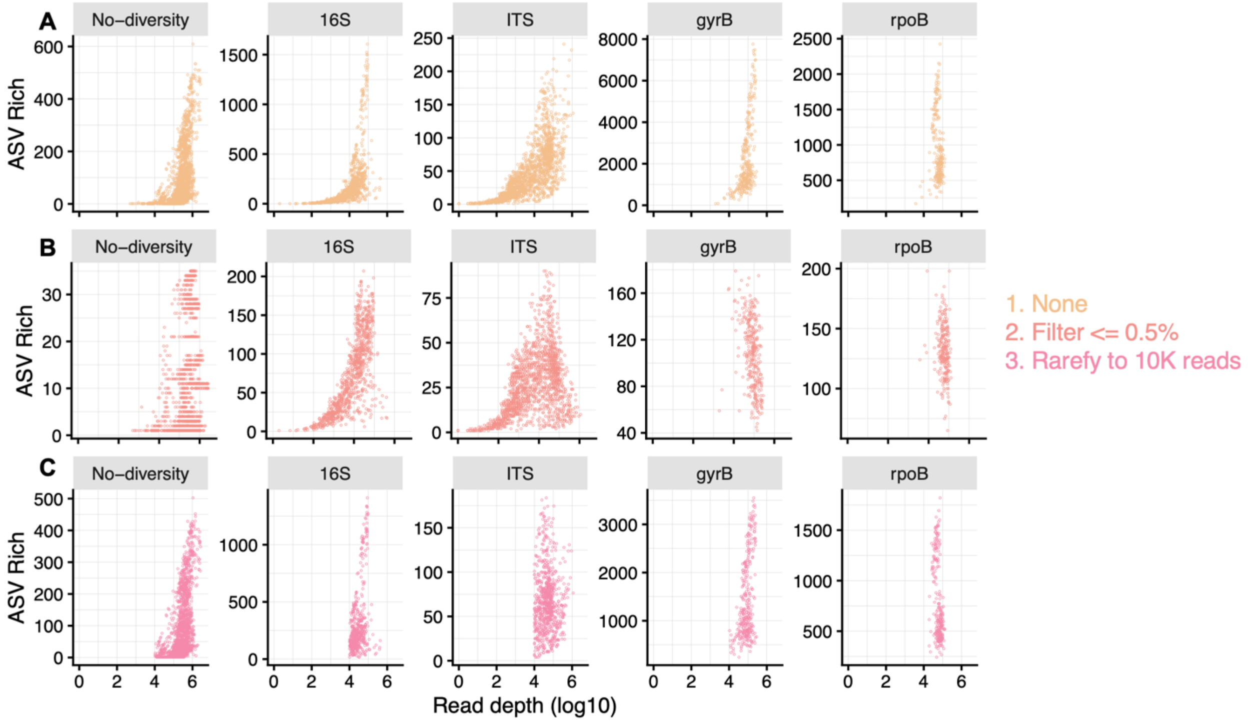


**Supp Fig 7:** Filtering and rarefying reduces the number of ASVs  but does not fix the dramatic increase in ASV richness above $10^4$ reads across amplicons. A) Unfiltered data, reproduced from Fig. 3. B) Filtering out ASVs at <=0.5% relative abundance reduces ASV richness at different magnitudes across amplicons, with a greater than 10-fold reduction for everything but ITS. However, the exponential increase above $10^4$ reads still exists for No-diversity, 16S, and ITS amplicons. C) Rarefying to 10,000 reads/sample reduces ASV richness less and obscures the exponential increase, but there are still hundreds of ASVs in No-diversity amplicons. Together, this suggests that filtering and rarefying alone are insufficient to resolve technical artefacts that impact microbial diversity estimates.

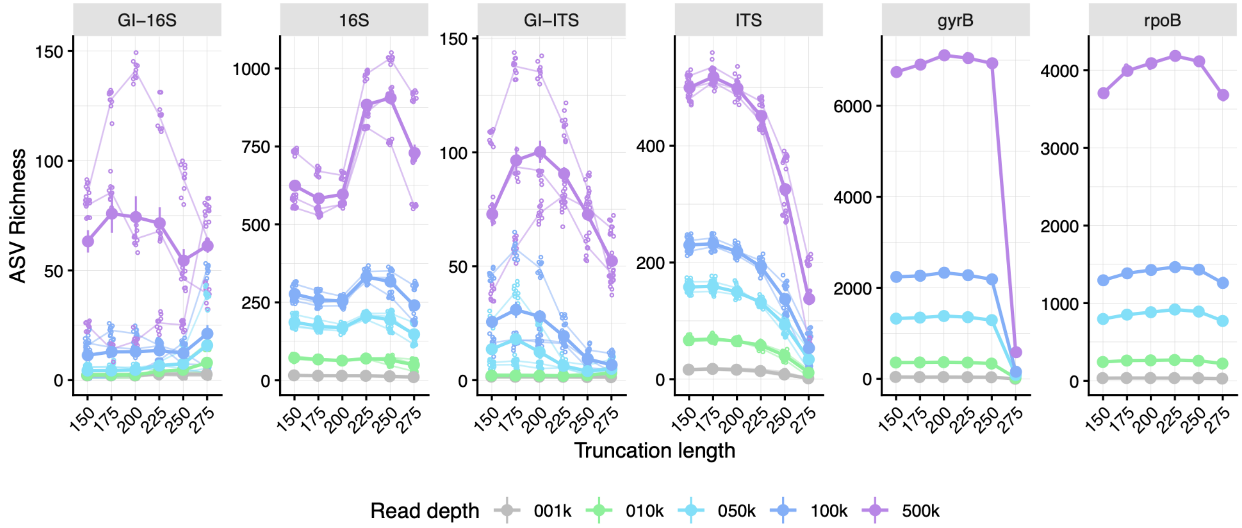


**Supp Fig 8:** Truncation length and read depth interacts to shape ASV richness across amplicons. Thick lines and points represent mean across sequencing runs. Thin lines show each run, and small points show the individual values (N=10/read depth); note that gyrB and rpoB were only sequenced on a single run.

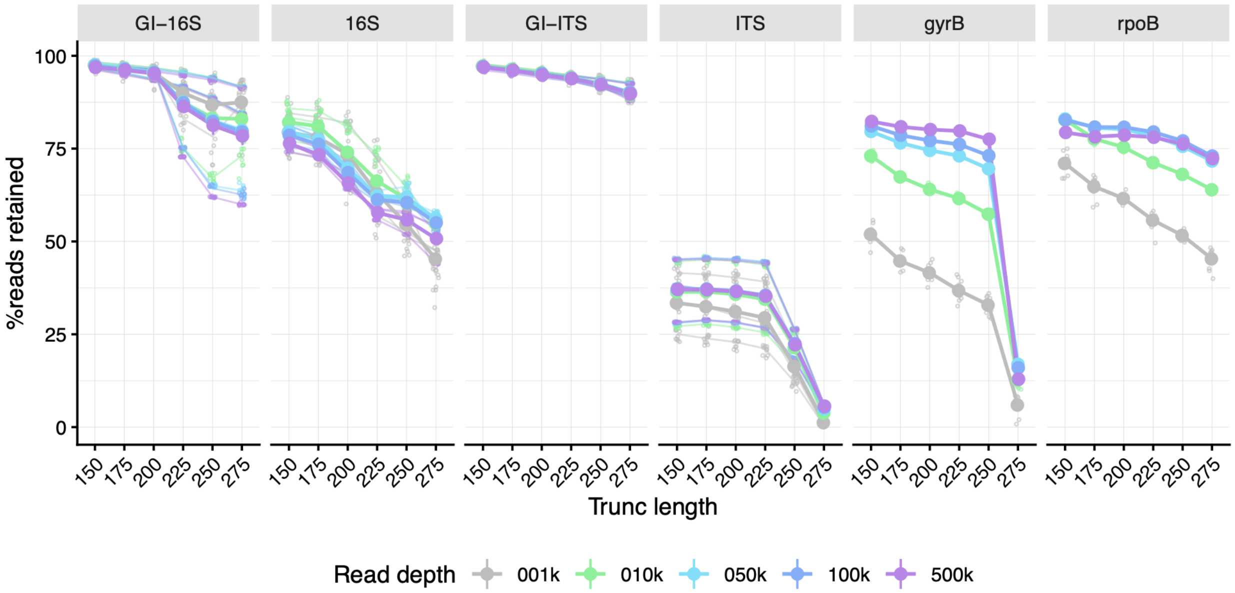


**Supp Fig 9:** Percentage of reads that remain after calling ASVs and assigning at least phylum-level taxonomy declines with longer truncation lengths. Thick lines and points show the average across all runs. Thin lines show each run, and small points show the individual values (N=10 per read depth); note that gyrB and rpoB were only sequenced on a single run. The magnitude of decline is different for each amplicon.

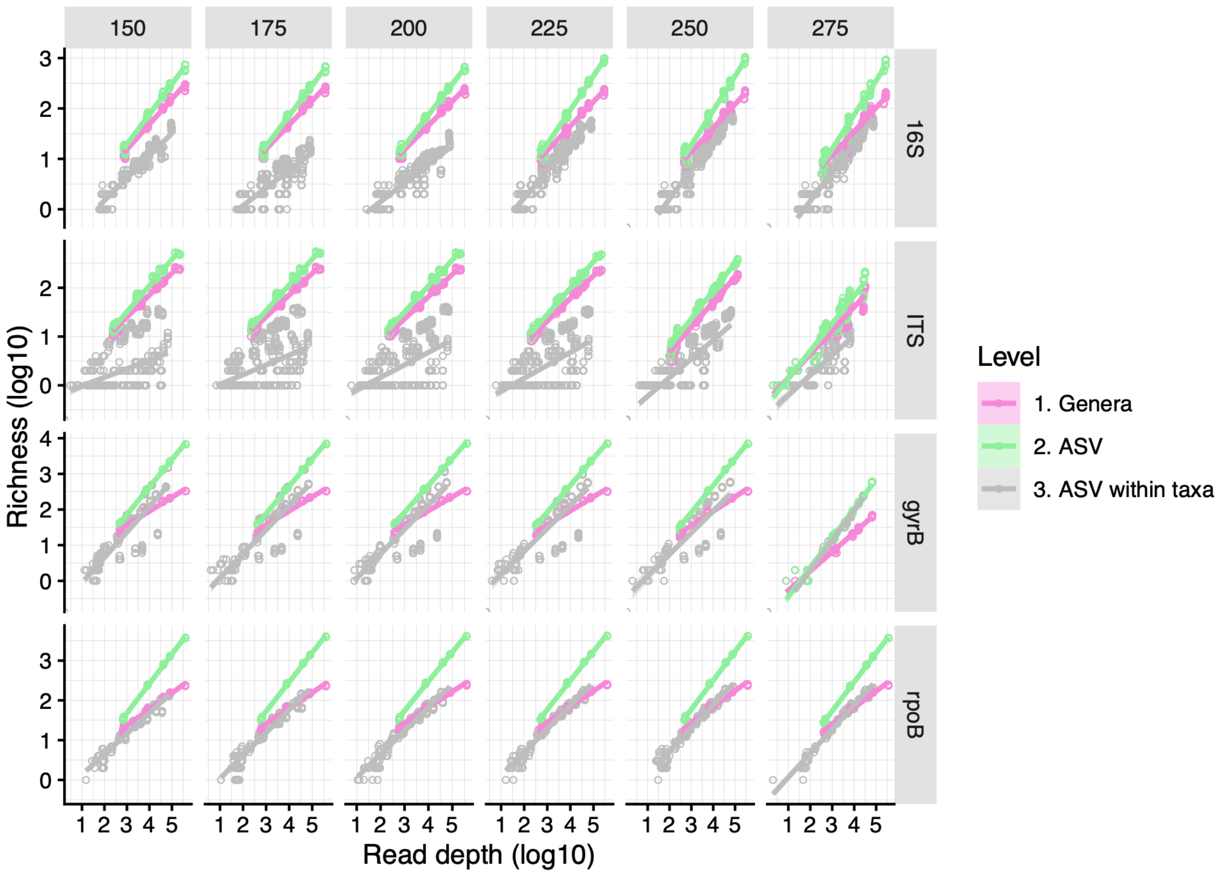


**Supp Fig 10:** Correlation between read depth and richness at either the genera level, ASV level, or ASVs within the top three most abundant taxa per amplicon. Facet columns are truncation lengths, and rows are amplicons. Each point represents the replicate per pooled subsampling depth at the different taxonomic levels. Note that for ITS and gyrB at 275 bp truncation length, >85% of reads were lost during ASV calling. The relationship between read depth and richness is shaped by significant interactions between trim length and taxonomic level (Wald interaction $X^2$ = 60.8711, df = 2, p < 0.0001, Supp. Table R6).

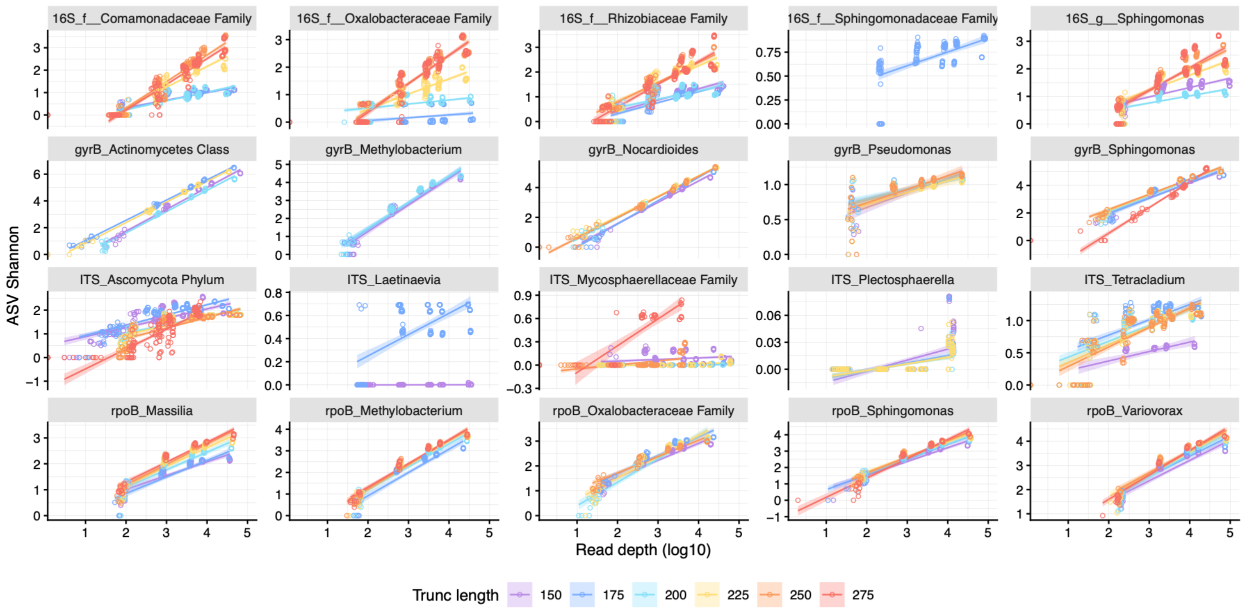


**Supp Fig 11:** Correlation between read depth and richness for the top 5 most abundant taxa assigned at the Genus level for each amplicon. Each point represents the Shannon diversity of ASVs within each taxa per replicate from read depth subsampling.

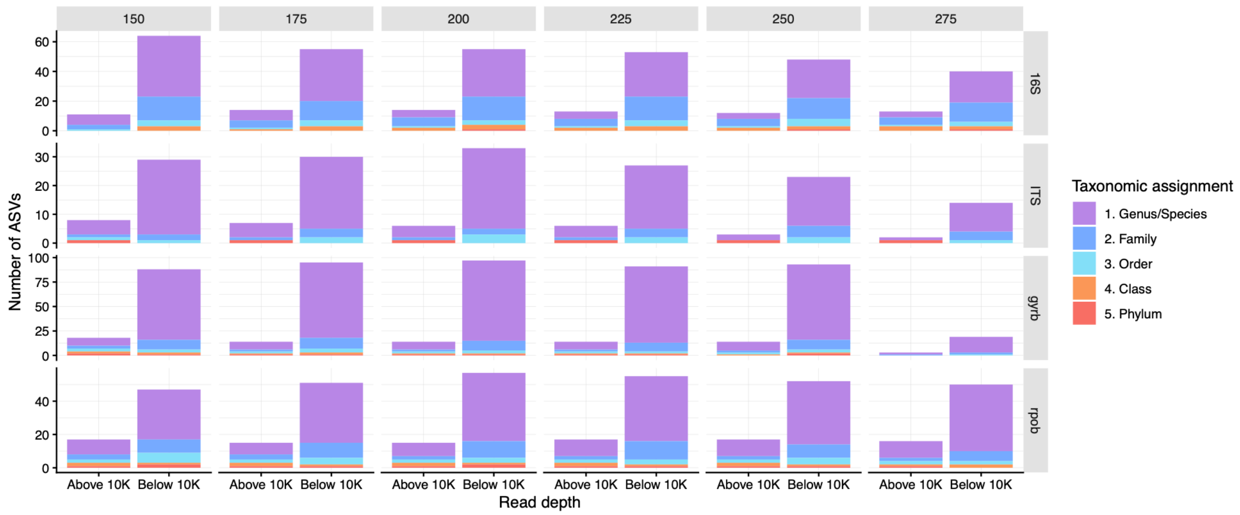


**Supp Fig 12:** Taxonomic assignment for taxa above $10^4$ or below $10^4$ reads, across all truncation lengths for each amplicon. Color represents the taxonomic assignment. Note that the y-axis is scaled per amplicon.

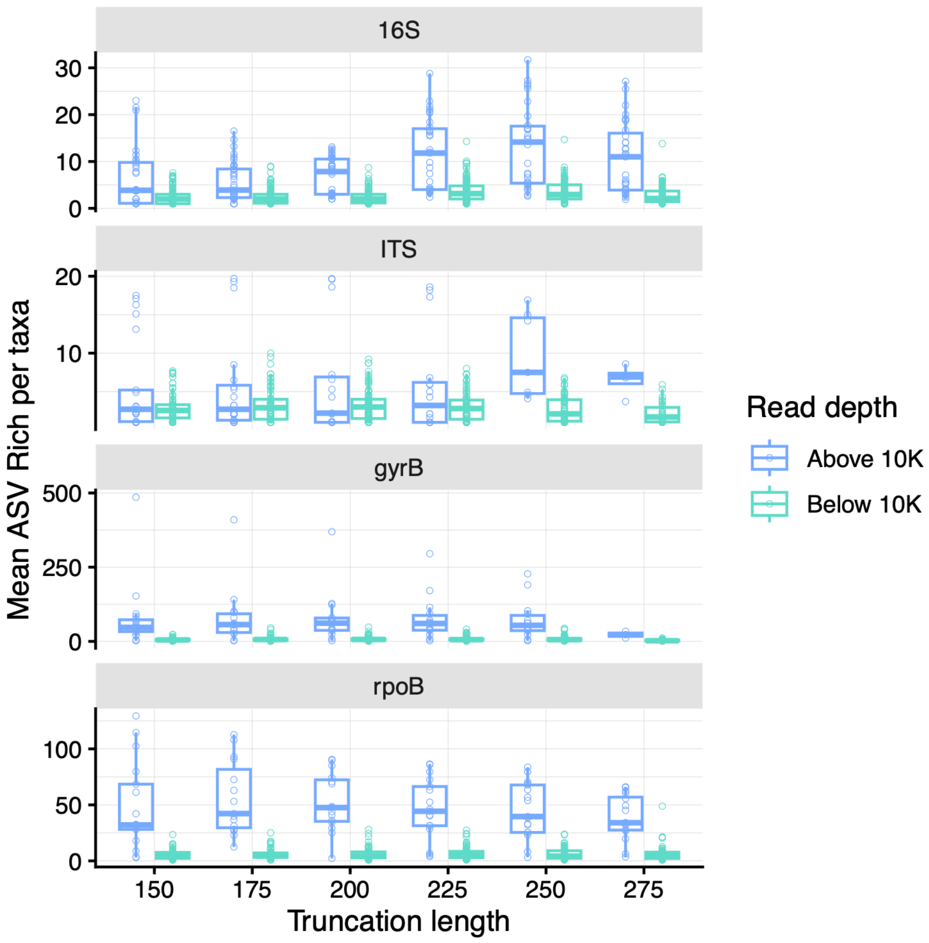


**Supp Fig 13:** Mean ASV richness within taxa, colored by whether the abundance of the taxa is above or below $10^4$ reads. Each point represents the ASV richness per taxa from each sequencing run across the four amplicons. The magnitude of increase in richness is more sensitive to read depth than for Shannon diversity (Fig. 4C).

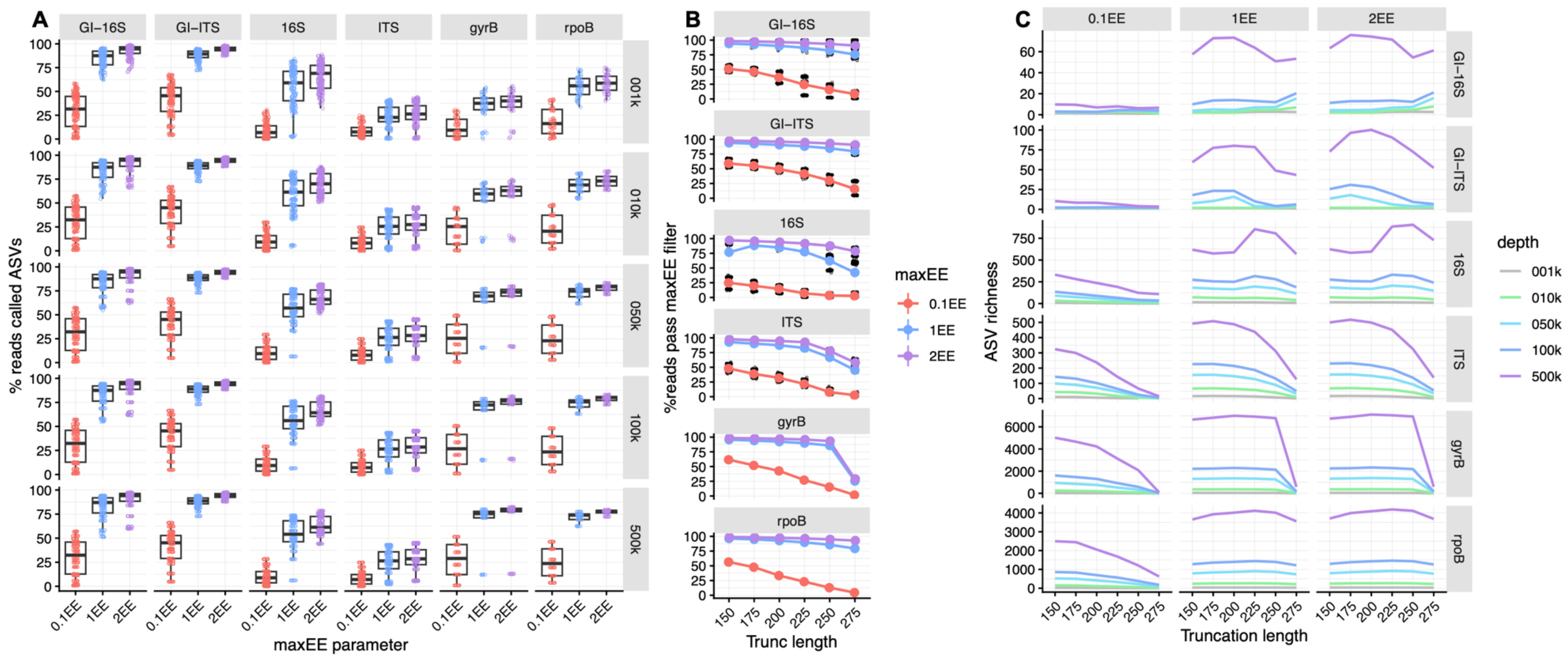


Supp Fig 14: Tuning maxEE parameter reduces read depth consistently across amplicons and removes spurious ASVs for GI only. A) Plots are faceted by amplicon and subsampled read depth. At a very strict maxEE = 0.1, ~75% of reads were removed—except for ITS, where all reads were removed. Moderate reductions were observed between maxEE = 2 and maxEE = 1. B) Percent reads passing filter is sensitive to truncation length but is overall much lower for maxEE =  0.1. Line represents the mean value for each maxEE parameter across truncation lengths. Small points represent individual samples. C) Plots are faceted by amplicon and maxEE value, and lines represent the average ASV richness across sequencing runs. maxEE = 0.1 succeeds in filtering many ASVs as compared to 1EE and 2EE values, but this is likely related to the substantial reduction in reads.

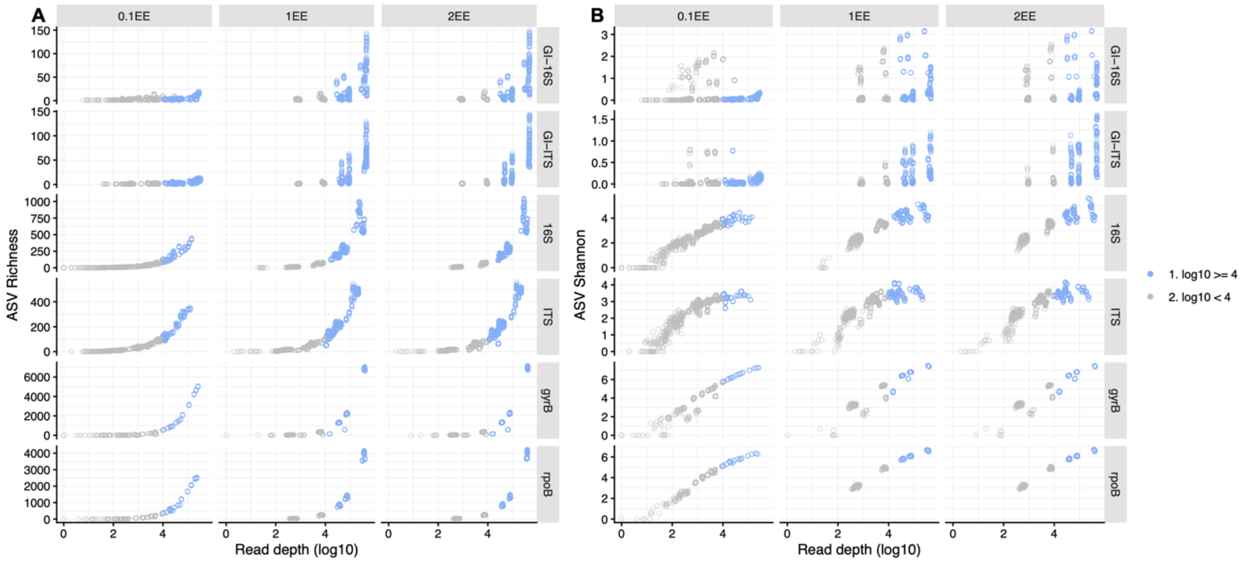


**Supp Fig 15:** Increasing sensitivity to max expected errors (maxEE) tempers the artificial inflation of diversity estimates above $10^4$ reads (blue points). Strictest filter is 0.1EE, while 2EE is least stringent and the default value in QIIME2. A) Strict maxEE = 0.1 filter succeeds in flattening the relationship between read depth and ASV richness for the no-diversity GI amplicon. The microbial amplicons still exhibit exponential increases, though the maximum is less compared to less stringent maxEE values. B) Shannon diversity responds similarly to strict maxEE filter, where no-diversity GI amplicons are essentially flattened, but the positive correlation remains for all other amplicons.

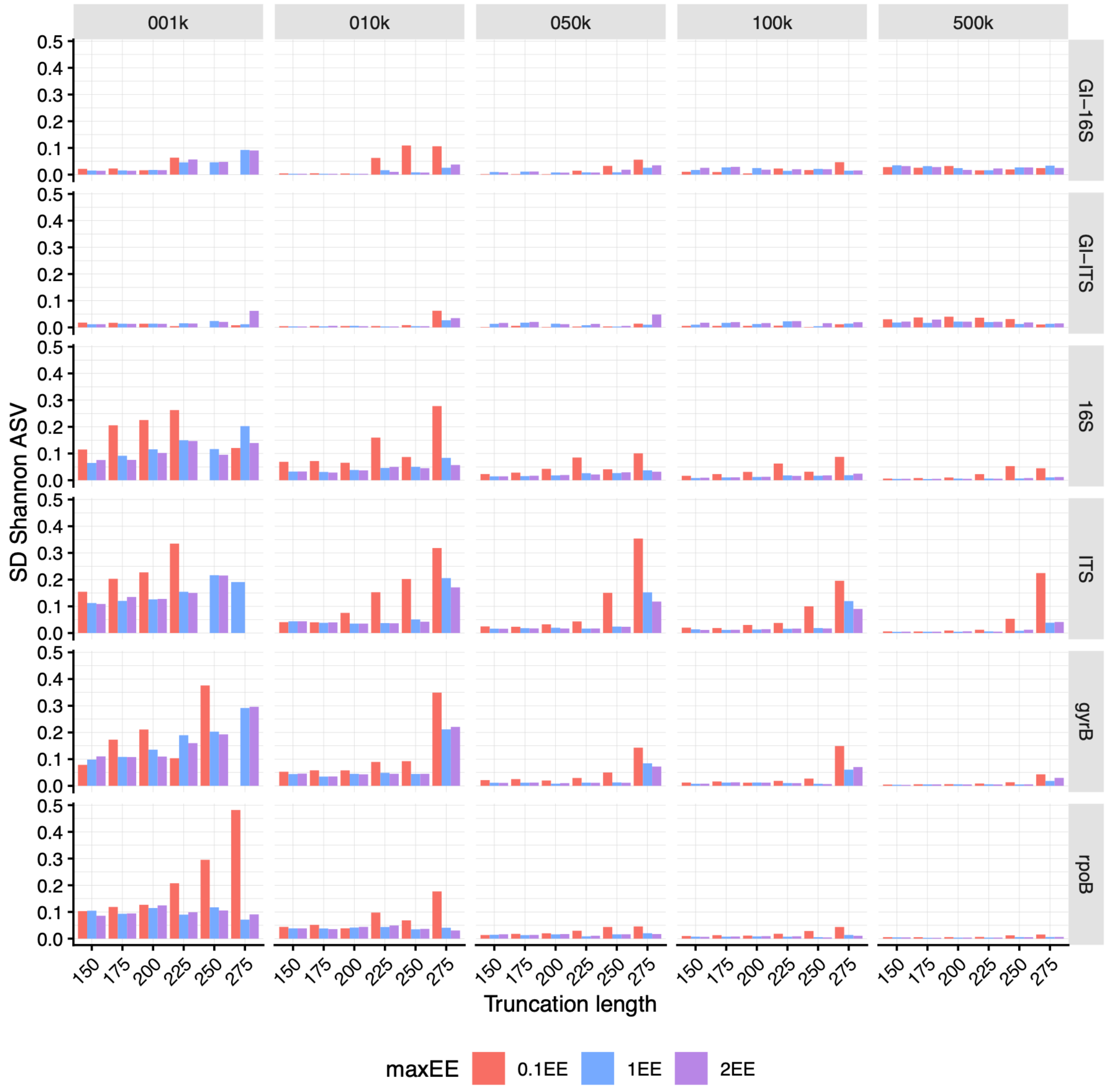


**Supp Fig 16:** Increasing sensitivity to max expected errors (maxEE) increases noise among samples in different ways across amplicons. Specifically, maxEE = 0.1 has high standard deviation at low read depths for all amplicons. For 16S and ITS, longer fragments over 200 bp have higher standard deviation at higher read depths, while gyrB and rpoB were more robust.

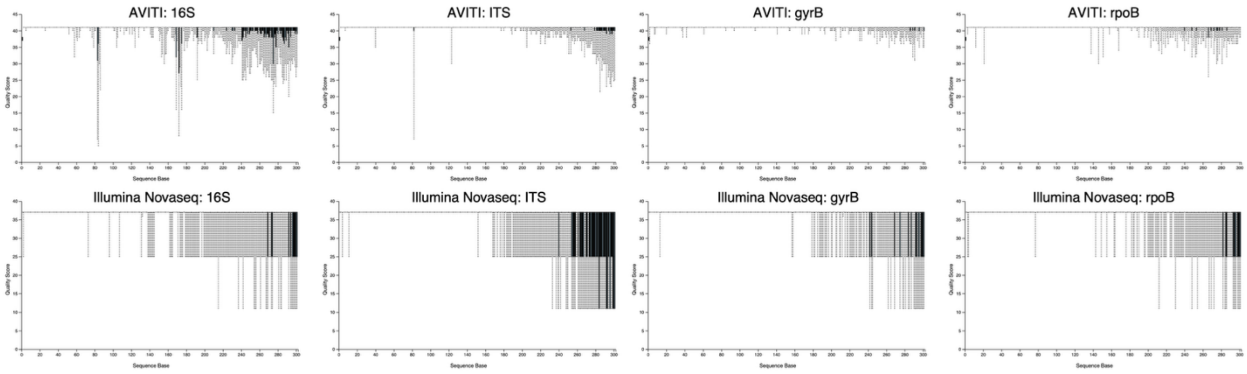


**Supp Fig 17:** Example traces for PHRED quality scores between samples sequenced on either AVITI or Novaseq platforms for the 16S, ITS, gyrB, and rpoB amplicons. Plots were generated using QIIIME2 demux visualization and are generated from the same library pools that were sequenced across the different platforms.  Note the difference in y-axis scales between AVITI and Novaseq, where AVITI tends to have PHRED scores above 40, while Illumina Novaseq have scores below 40.

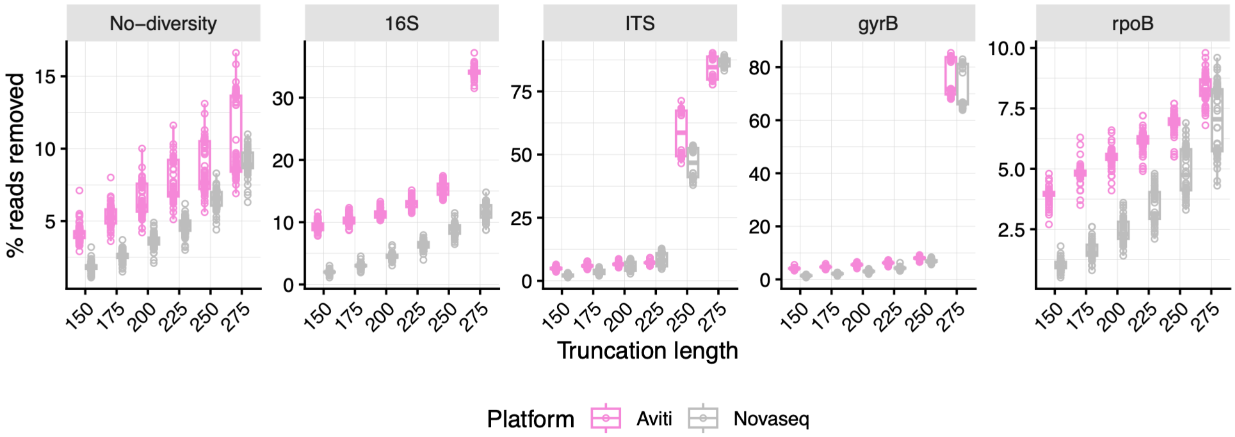


**Supp Fig 18:** More reads are removed during the initial maxEE filtering in AVITI than Novaseq data across all amplicons and truncation lengths. Each point represents a replicate from the subsampling at different depths (i.e., 1k, 10k, 50k, 100k, 500k) from plant and soil samples. Note that the “No-diversity” amplicons include GI co-amplified with either 16S or ITS from plant samples or the 16S spike-in amplified in soil samples.

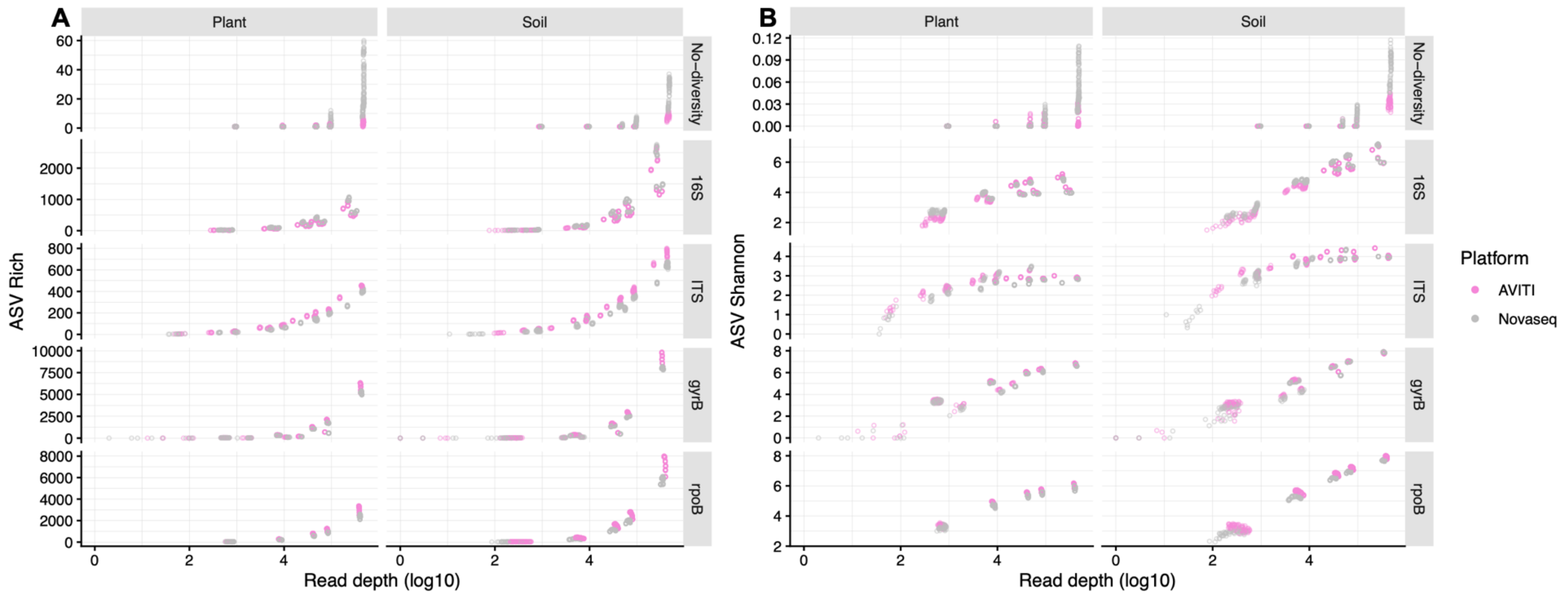


**Supp Fig 19:** ASV diversity estimates are impacted by read depth across both AVITI and Novaseq platforms, but the sequencing platform differentially impacts the magnitude and directionality of impact across amplicons (Supp. Table R11). Points represent the replicates at each subsampled read depth for the six different truncation lengths. Note that the "No-diversity" for the plant facet is GI ASVs co-amplified with either 16S or ITS, but 16S spike-in ASVs for the soil facet. Also, for ITS and gyrB, like in previous analyses, truncation length of 275 bp results in > 80% read loss. A) Exponential increases in ASV richness were still detected at above $10^4$ reads. For only the no-diversity and 16S amplicons, ASV richness was higher for data generated using the Novaseq platform. ITS, gyrB, and rpoB displayed higher ASV richness for data generated using the AVITI platform. B) Shannon diversity also increased with read depth, though the effects were tempered compared to ASV richness.

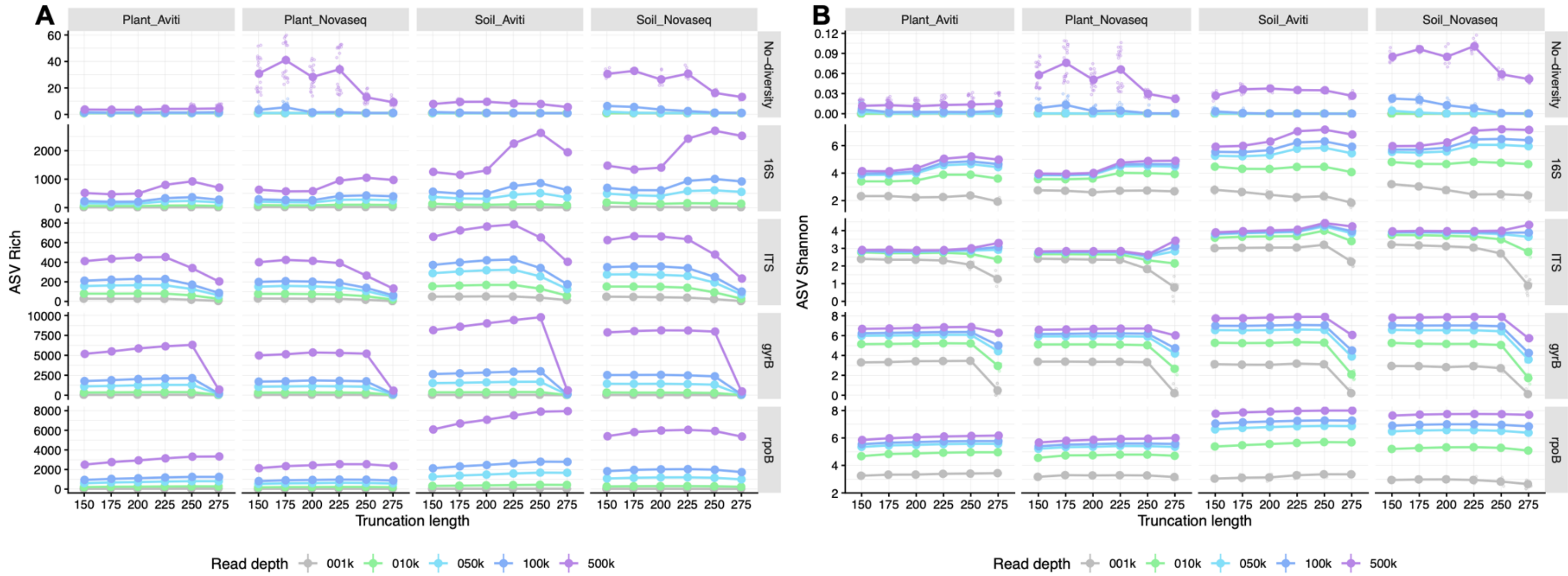


**Supp Fig 20:** AVITI and Novaseq sequencing platforms both show impacts of truncation length and read depth on ASV Shannon diversity. Thick lines and points represent mean ± standard error across all sequencing runs per sequencing depth. Plots are faceted by plant and soil per sequencing platform. Note that the “No-diversity” for the plant facet is GI ASVs co-amplified with either 16S or ITS, but 16S spike-in ASVs for the soil facet. Also, for ITS and gyrB, like in previous analyses, truncation length of 275 bp results in > 80% read loss. A) ASV richness estimates are higher for No-diversity amplicons in the Novaseq sequencing compared to AVITI sequencing. 16S and and ITS are similar between the two platforms. ASV richness is higher in AVITI sequencing for the high-resolution gyrB and rpoB amplicons. The GI amplicons display a distinct interaction between truncation length and high sequencing depth (500k reads/sample), but otherwise the effects of truncation length are similar across amplicons. B) Similar dynamics for ASV Shannon diversity were observed as for ASV richness.

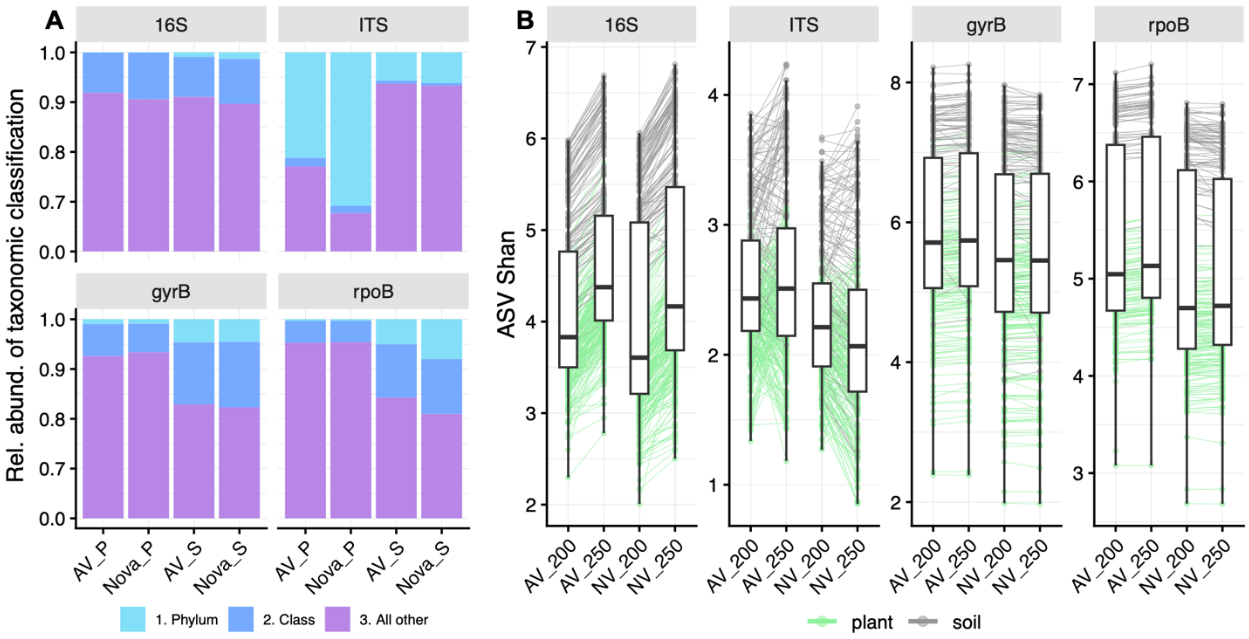


**Supp. Fig. 21**: A) Bars represent the ASV taxonomic level assignment for either AVITI (AV) or Novaseq (Nova) for plant (P) and soil (S) samples. Higher percentage of the microbiome is unresolved beyond phylum or class taxonomic levels for data generated on the Novaseq platform than on AVITI. Note the y-axis break between 0.0 and 0.70 to facilitate visualization. B) Differences in Shannon diversity between 200 and 250 bp truncation lengths, grouped by sequencing platform. Color represents the sample type, and lines connect individual samples between truncation lengths. ITS data were particularly inconsistent between truncation lengths, which is likely related to inconsistencies in taxonomic assignment shown in panel A.

Supp. Table 1: Kruskal-Wallis test values for comparing mean GI Shannon diversity between runs at two truncation lengths.

| | Kruskal-Wallis $X^2$ | df | p |
|---|---|---|---|
| 200 bp | 5.3524 | 1 | 0.02* |
| 250 bp | 54.76 | 1 | <0.0001*** |

Supp. Table 2: Model fits for comparing correlation between 200 and 250 bp for raw, filtered (<0.5% relative abundance removed), and rarefied (10,000 reads/sample).

| Trunc. | Modification | adj. $R^2$ | Test stat | df | P-value |
|---|---|---|---|---|---|
| 200 bp | 1. none | 0.6919167 | 48.1633826 | 1 | <0.0001*** |
| 250 bp | 1. none | 0.37952041 | 13.233131 | 1 | 0.002** |
| 200 bp | 3. rarefy | 0.6445355 | 39.077629 | 1 | <0.0001*** |
| 250 bp | 3. rarefy | 0.27320279 | 8.51799236 | 1 | 0.009** |
| 200 bp | 2. filter | 0.52957185 | 24.6401859 | 1 | <0.0001*** |
| 250 bp | 2. filter | 0.03250439 | 1.67192852 | 1 | 0.212 |

Supp. Table 3: Kruskal-Wallis test values for comparing ASV richness between raw and rarefied (10,000 reads/sample) for the five different amplicons.

| Amplicon | Kruskal-Wallis $X^2$ | df | p-value |
|---|---|---|---|
| No-diversity | 0.82 | 1 | 0.3653 |
| 16S | 0.62 | 1 | 0.4299 |
| ITS | 24.31 | 1 | <0.0001*** |
| gyrB | 39.92 | 1 | <0.0001*** |
| rpoB | 26.32 | 1 | <0.0001*** |

Supp. Table 4: Fixed effects for the analysis of read depth and truncation length on Shannon diversity. Significance was evaluated with Type III Wald $X^2$ tests with Kenward-Roger degrees of freedom.

| Model: Shannon ~ truncation length * amplicon * log10(read depth) + (1\| seq. run) | | | |
|---|---|---|---|
| | $X^2$ | Df | P-value |
| (Intercept) | 27.271 | 1 | 1.77E-07*** |
| Truncation length | 248.15 | 5 | < 2.2e-16*** |
| Amplicon | 142.476 | 5 | < 2.2e-16*** |
| Log10(read depth) | 333.442 | 1 | < 2.2e-16*** |
| Trunc. length * amplicon | 751.84 | 25 | < 2.2e-16*** |
| Trunc. Length * log10(read depth) | 368.98 | 5 | < 2.2e-16*** |
| Amplicon * log10(read depth) | 702.543 | 5 | < 2.2e-16*** |
| Trunc. Length * amplicon * log10(read depth) | 531.077 | 25 | < 2.2e-16*** |

Supp. Table 5: Fixed effects for the analysis of read depth and truncation length on ASV richness (log10-transformed). Significance was evaluated with Type III Wald $X^2$ tests with Kenward-Roger degrees of freedom.

| Model: ASV rich ~ truncation length * amplicon * log10(read depth) + (1\| seq. run) | | | |
|---|---|---|---|
| | $X^2$ | Df | P-value |
| (Intercept) | 29.587 | 1 | 5.35E-08*** |
| Truncation length | 27.557 | 5 | 4.44E-05*** |
| Amplicon | 220.19 | 5 | < 2.2e-16*** |
| Log10(read depth) | 861.169 | 1 | < 2.2e-16*** |
| Trunc. length * amplicon | 276.916 | 25 | < 2.2e-16*** |
| Trunc. Length * log10(read depth) | 26.901 | 5 | 5.96E-05*** |
| Amplicon * log10(read depth) | 101.667 | 5 | < 2.2e-16*** |
| Trunc. Length * amplicon * log10(read depth) | 131.841 | 25 | < 2.2e-16*** |

Supp. Table 6: Fixed effects for the analysis of read depth, truncation length, and taxonomic level (ASV, species, genus) on ASV richness (log10-transformed). Significance was evaluated with Type III Wald $X^2$ tests with Kenward-Roger degrees of freedom.

| Model: ASV rich ~ log10(read depth) * trunc. length * tax. level + (1\| amplicon) + 1(seq. run) | | | |
|---|---|---|---|
| | $X^2$ | Df | P-value |
| (Intercept) | 7.0486 | 1 | 7.93E-03** |
| log10(read depth) | 99.9902 | 1 | < 2.2e-16*** |
| Truncation length | 8.2058 | 1 | 0.004176** |
| Taxonomic level | 9.8036 | 2 | 7.43E-03** |
| log10(read depth) * trunc. length | 1.5846 | 1 | 0.208093 |
| log10(read depth) * taxonomic level | 95.9493 | 2 | < 2.2e-16*** |
| Trunc. Length * taxonomic level | 1.1142 | 2 | 0.572872 |
| log10(read depth) * trunc. length. * tax. level | 60.8711 | 2 | 6.05E-14*** |

Supp. Table 7: Fixed effects for the analysis of read depth, truncation length, and taxonomic level (ASV, species, genus) on ASV Shannon diversity within individual taxa (square root-transformed). Significance was evaluated with Type III Wald $X^2$ tests with Kenward-Roger degrees of freedom.

| Model: ASV Shannon ~ log10(read depth) * trunc. length * amplicon + 1(seq. run) | | | |
|---|---|---|---|
| | $X^2$ | Df | P-value |
| (Intercept) | 84.7238 | 1 | < 2.2e-16*** |
| log10(read depth) | 20.2127 | 1 | 6.93E-06*** |
| Truncation length | 0.8172 | 3 | 0.84534 |
| Amplicon | 10.2234 | 1 | 0.001387** |
| log10(read depth) * amplicon | 22.1581 | 3 | 6.05E-05*** |
| log10(read depth) * trunc. length | 63.1681 | 1 | 1.90E-15*** |
| Amplicon * trunc. length | 2.4231 | 3 | 0.489344 |
| log10(read depth) * amplicon * trunc. length | 11.8064 | 3 | 0.008077** |

Supp. Table 8: Fixed effects for the analysis of taxonomic level and truncation length on ASV richness differences between abundant taxa (>= 10000 reads) or rare taxa (>1000 reads, < 10000 reads). Significance was evaluated with Type II Wald $X^2$ tests with Kenward-Roger degrees of freedom.

| Model: log10(ASV rich) ~ tax. level * truncation length + (1\|amplicon) + (1\|seq. run) | | | |
|---|---|---|---|
| | $X^2$ | Df | Pr(>Chisq) |
| Taxonomic level | 934.474 | 1 | < 2.2e-16*** |
| Truncation length | 14.783 | 1 | 0.0001206*** |
| Trunc. length * taxonomic lev. | 13.066 | 1 | 0.0003008*** |

Supp. Table 9: Fixed effects for the analysis of taxonomic level and truncation length on ASV Shannon diversity differences between abundant taxa (>= 10000 reads) or rare taxa (>1000 reads, < 10000 reads). Significance was evaluated with Type II Wald $X^2$ tests with Kenward-Roger degrees of freedom.

| Model: log10(ASV rich) ~ tax. level * truncation length + (1\|amplicon) + (1\|seq. run) | | | |
|---|---|---|---|
| | $X^2$ | Df | Pr(>Chisq) |
| Taxonomic level | 934.474 | 1 | < 2.2e-16*** |
| Truncation length | 14.783 | 1 | 0.0001206*** |
| Trunc. length * taxonomic lev. | 13.066 | 1 | 0.0003008*** |

Supp. Table 10: Fixed effects for the analysis of maxEE parameter on noise in Shannon diversity measurements. Noise was estimated as the square-root transformed, standard deviation across replicates following different maxEE trim parameters. Significance was evaluated with Type III Wald $X^2$ tests with Kenward-Roger degrees of freedom

| Model: sd(Shannon) ~ maxEE * trunc. length * log10(read depth) + (1\|amplicon) + (1\| seq.run) | | | |
|---|---|---|---|
| | $X^2$ | Df | P-value |
| (Intercept) | 97.183 | 1 | < 2.2e-16*** |
| maxEE parameter | 12.177 | 2 | 0.0022684** |
| Truncation length | 15.21 | 1 | 9.62E-05*** |
| log10(read depth) | 64.404 | 1 | 1.01E-15*** |
| maxEE * trunc. length | 15.663 | 2 | 0.000397*** |
| maxEE * log10(read depth) | 13.513 | 2 | 0.0011634** |
| Trunc. length * log10(read depth) | 19.82 | 1 | 8.51E-06*** |
| maxEE * trunc. length * log10(read depth) | 15.987 | 2 | 0.0003376*** |

Supp. Table 11: Fixed effects for the analysis of sequencing platform (AVITI Element versus Illumina Novaseq) on ASV richness. Significance was evaluated with Type II Wald $X^2$ tests with Kenward-Roger degrees of freedom.

| Model: log10(ASV rich) ~ platform * trunc. length + log10(read depth) + (1\|amplicon) | | | |
|---|---|---|---|
| | $X^2$ | Df | P-value |
| Platform | 28.3169 | 1 | 1.03E-07*** |
| Truncation length | 4.6035 | 1 | 0.03191* |
| log10(read depth) | 451.3937 | 1 | < 2.2e-16*** |
| Platform * trunc. length | 3.9405 | 1 | 0.04714* |

Supp. Table 12: Fixed effects for the analysis of sequencing platform on the percentage of reads removed (log10-transformed) during the initial filtering step of the DADA2 algorithm. Random effects controlled for truncation length and sample type (soil versus plant). Significance was evaluated with Type II Wald $X^2$ tests with Kenward-Roger degrees of freedom.

| Model: log10(% reads remove) ~ platform * amplicon + log10(reads) + (1\|trunc. length) + (1\|sample type) | | | |
|---|---|---|---|
| | $X^2$ | Df | P-value |
| Platform | 2821.454 | 1 | < 2.2e-16*** |
| Amplicon | 4228.548 | 4 | < 2.2e-16*** |
| log10(read depth) | 91.343 | 1 | < 2.2e-16*** |
| Platform * amplicon | 469.059 | 4 | < 2.2e-16*** |

Supp. Table 13: Fixed effects for the analysis of sequencing platform, read depth, amplicon, and truncation length on either ASV richness or ASV Shannon diversity. ASV richness was log10-transformed, and ASV Shannon was square root-transformed. Four-way interactions were tested for overfitting by comparing AIC with reduced models. Significance was evaluated with Type III Wald $X^2$ tests with Kenward-Roger degrees of freedom.

| Model: ASV div ~ log10(reads) * platform * amplicon * trunc. length + (1\| sample.type) | | | | | |
|---|---|---|---|---|---|
| | Df | Rich $X^2$ | Rich P-val | Shan $X^2$ | Shan P-val |
| (Intercept) | 1 | 20.24 | 6.83E-06*** | 1.0492 | 0.3057 |
| log10(read depth) | 1 | 48.386 | 3.50E-12*** | 5.097 | 0.0240* |
| Platform | 1 | 82.877 | < 2.2e-16*** | 8.1747 | 0.0042** |
| Amplicon | 4 | 96.924 | < 2.2e-16*** | 580.269 | < 2.2e-16*** |
| Truncation length | 1 | 0.518 | 4.72E-01 | 0.0222 | 0.8814 |
| log10(reads) * platform | 1 | 138.681 | < 2.2e-16*** | 13.4659 | 0.0002*** |
| log10(reads) * amplicon | 4 | 163.472 | < 2.2e-16 | 234.9685 | < 2.2e-16*** |
| Platform * amplicon | 4 | 84.535 | < 2.2e-16 | 41.5928 | 2.03E-08*** |
| log10(reads) * trunc. length | 1 | 0.876 | 0.3493 | 0.0617 | 0.8038 |
| Platform * trunc. length | 1 | 31.236 | 2.29E-08*** | 3.4 | 0.0652 |
| Amplicon * trunc. length | 4 | 62.391 | 9.12E-13*** | 375.4082 | < 2.2e-16*** |
| reads * platform * amplicon | 4 | 117.166 | < 2.2e-16*** | 42.0459 | 1.63E-08*** |
| reads * platform * trunc. len. | 1 | 55.188 | 1.10E-13*** | 5.8274 | 0.01578* |
| reads * amplicon * trunc. len | 4 | 42.502 | 1.31E-08*** | 328.21 | < 2.2e-16*** |
| Platform * amplicon * trunc. len | 4 | 40.643 | 3.19E-08*** | 46.6152 | 1.83E-09*** |
| 4-way interaction | 4 | 48.931 | 6.04E-10*** | 39.6504 | 5.11E-08*** |

Supp. Table 14: Fixed effects for the analysis of sequencing platform, sample type (soil versus plant) and amplicon on ASV richness and Shannon diversity estimates from true samples. True samples were normalized to same depth prior to DADA2 ASV calling. For both ASV richness and Shannon diversity, the four-way interaction was initially fit as above (e.g., Supp. Table 13), but was not significant. ASV richness was log10-transformed, and ASV Shannon was square root-transformed. The subsequent model only included significant three- and two-way interactions. The models were tested for overfitting by comparing AIC with reduced models. Significance was evaluated with Type III Wald $X^2$ tests with Kenward-Roger degrees of freedom.

| Model: Diversity ~ platform * sample type * amp. + trunc. length * sample type *amplicon + log10(read depth) + (1\|DNA plate) | | | | | |
|---|---|---|---|---|---|
| | Df | Rich $X^2$ | Rich p-val | Shan $X^2$ | Shan p-val |
| (Intercept) | 1 | 43.2893 | 4.72E-11*** | 1976.4092 | < 2.2e-16*** |
| Platform | 1 | 41.9995 | 9.13E-11*** | 63.8683 | 1.33E-15*** |
| Sample type | 1 | 978.9796 | < 2.2e-16*** | 658.9971 | < 2.2e-16*** |
| Amplicon | 3 | 1197.591 | < 2.2e-16*** | 353.3264 | < 2.2e-16*** |
| Truncation length | 1 | 730.9918 | < 2.2e-16*** | 169.7177 | < 2.2e-16*** |
| log10(read depth) | 1 | 3096.6778 | < 2.2e-16*** | 80.7335 | < 2.2e-16*** |
| Platform * sample type | 1 | 38.622 | 5.14E-10*** | 69.6168 | < 2.2e-16*** |
| Platform * amplicon | 3 | 490.7545 | < 2.2e-16*** | 8.4228 | 0.038035* |
| Sample type * amplicon | 3 | 38.6609 | 2.05E-08*** | 76.2437 | < 2.2e-16*** |
| Sample type * trunc. length | 1 | 5.7531 | 0.01646* | 3.0137 | 0.082562 |
| Amplicon * trunc. length | 3 | 381.9039 | < 2.2e-16*** | 195.1132 | < 2.2e-16*** |
| Platform * sample type * amp. | 3 | 15.8325 | 0.001227** | 59.4424 | 7.73E-13*** |
| Sample.type * amp. * trunc. len | 3 | 26.1905 | 8.70E-06*** | 13.0863 | 0.004454** |

Supp. Table 15: Adjusted $R^2$ values for the comparison between 200 and 250 bp truncation length per each platform and amplicon.

| Amplicon | Sample type | Measure | AVITI $R^2$ | Novaseq $R^2$ |
|---|---|---|---|---|
| 16S | plant | ASV Rich | 0.979 | 0.956 |
| 16S | soil | ASV Rich | 0.988 | 0.962 |
| 16S | plant | ASV Shannon | 0.8 | 0.813 |
| 16S | soil | ASV Shannon | 0.978 | 0.898 |
| gyrB | plant | ASV Rich | 0.999 | 0.983 |
| gyrB | soil | ASV Rich | 0.999 | 0.971 |
| gyrB | plant | ASV Shannon | 0.997 | 0.994 |
| gyrB | soil | ASV Shannon | 0.98 | 0.978 |
| ITS | plant | ASV Rich | 0.809 | 0.729 |
| ITS | soil | ASV Rich | 0.762 | 0.813 |
| ITS | plant | ASV Shannon | 0.0229 | 0.36 |
| ITS | soil | ASV Shannon | 0.404 | 0.802 |
| rpoB | plant | ASV Rich | 0.996 | 0.969 |
| rpoB | soil | ASV Rich | 0.996 | 0.897 |
| rpoB | plant | ASV Shannon | 0.997 | 0.996 |
| rpoB | soil | ASV Shannon | 0.996 | 0.945 |

Supp. Table 16: Summary of sample types and platforms in the eight different sequencing runs.

| Sequencing run | Platform | Samples | Amplicons |
|---|---|---|---|
| Run01 | Illumina Novaseq | Michigan | GI, 16S |
| Run02 | Illumina Novaseq | Michigan | GI, 16S |
| Run03 | Illumina Novaseq | Michigan | GI, 16S |
| Run04 | Illumina Novaseq | Michigan | GI, ITS |
| Run05 | Illumina Novaseq | Michigan | GI, ITS |
| Run06 | Illumina Novaseq | Michigan | GI, ITS |
| Run07 | Illumina Novaseq | NYC | GI, 16S, ITS, gyrB, rpoB |
| Run08 | AVITI Element | NYC | GI, 16S, ITS, gyrB, rpoB |

Supp. Table 17: PCR conditions for each of the amplicons.

| Amplicon | Anneal temp | $1^{st}$ PCR cycles |
|---|---|---|
| 16S | 59ºC | 25 |
| ITS | 59ºC | 25 |
| gyrB | 68ºC | 30 |
| rpoB | 69ºC | 30 |
